# Venomics and Peptidomics of Palearctic vipers: Clade-wide analysis of seven taxa of the genera *Vipera*, *Montivipera*, *Macrovipera* and *Daboia* across Türkiye

**DOI:** 10.1101/2024.03.04.583389

**Authors:** Maik Damm, Mert Karış, Daniel Petras, Ayse Nalbantsoy, Bayram Göçmen, Roderich D. Süssmuth

## Abstract

Snake venom variations are a crucial factor to understand the consequences of snakebite envenoming worldwide and therefore it’s important to know about toxin composition alterations between taxa. Palearctic vipers of the genera *Vipera*, *Montivipera*, *Macrovipera* and *Daboia* have high medical impacts across the Old World. One hotspot for their occurrence and diversity is Türkiye on the border between the continents, but many of their venoms remain still understudied. Here, we present the venom compositions of seven Turkish viper taxa. By complementary mass spectrometry-based bottom-up and top-down workflows, the venom profiles were investigated on proteomics and peptidomics level. This study includes the first venom descriptions of *Vipera berus barani*, *Vipera darevskii*, *Montivipera bulgardaghica albizona* and *Montivipera xanthina*, as well as first snake venomics profiles of Turkish *Macrovipera lebetinus obtusa* and *Daboia palaestinae*, including an in-depth reanalysis of *Montivipera bulgardaghica bulgardaghica* venom. Additionally, we identified the modular consensus sequence pEXW(PZ^1–2P^(EI)/(KV)PPLE for bradykinin-potentiating peptides (BPP) in viper venoms. For better insights into variations and potential impacts of medical significance the venoms were compared against other Palearctic viper proteomes, including the first genus-wide *Montivipera* venom comparison. This will help the risk assessment of snakebite envenoming by these vipers and aid in predicting the venoms pathophysiology and clinical treatments.

## 1. INTRODUCTION

Snakebite envenoming is a major burden on global health ^1–3^. More than 5.4 million annual snakebites cause more than 150,000 casualties and several more long-lasting physical as well as often neglected mental disabilities ^4–7^. Responsible for a high number of these snake encounters are, beside elapids (Elapidae) and pit vipers (Crotalinae), the “true” or Old World vipers (Viperinae) ^8^. The occurrence of Old World vipers is distributed from the European Atlantic coast across the Palearctic realm, North Africa, Arabia peninsular to Asia and the Pacific coast in the Far East ^9–11^. Several taxa within this subfamily are in the focus of epidemiological snakebite envenoming dynamics and venom research ^12–17^. Among them, are the particularly relevant Palearctic vipers of the genera: *Vipera*, *Montivipera*, *Macrovipera* and *Daboia*, following Freitas *et al*. 2020. They consist of about 35 species, but their taxonomic classification has been a topic of debate for long time ^10,18,19^. The World Health Organization WHO lists all four genera at the highest medical importance, Category 1, with strong impact across their distributions ^8,11,13,20–22^.

Viper envenomation are characterized by mostly hemotoxic and tissue damaging clinical effects, while several Viperinae venoms, such as from the Russell’s viper *Daboia russelii* or the nose-horned viper *Vipera ammodytes* are also known for their capability to cause neurotoxic effects ^23–26^. Responsible for this spectrum of symptoms are more than 50 known toxin families in snake venoms, which are physiologically diverse: they occur in multiple isoforms and are functionally modulated via posttranslational modifications ^27–29^. Viperine venoms are primarily composed by enzymatic (e.g. proteases, lipases, oxidases) and non-enzymatic (e.g. lectins, growth factors, hormones) components extending molecular sizes across four magnitudes from small peptides of <500 Da up to protein complexes of >120 kDa ^30,31^. Over the last decade, venoms of Palearctic vipers have been intensively analysed on the proteomic level for 20 species across 25 countries (**Figure 1**).

**Figure 1.**
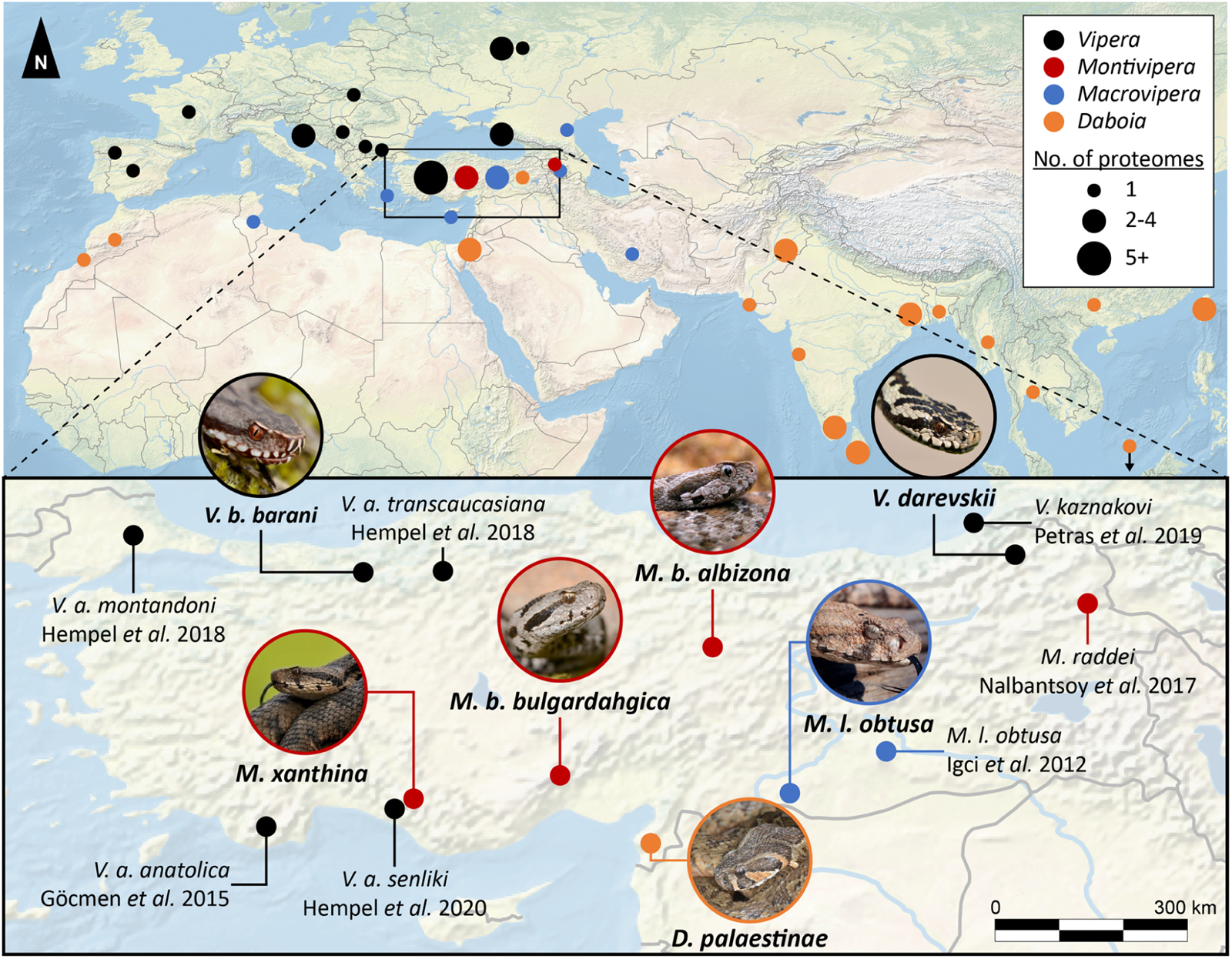
Mapped venomics studies of four Palearctic viper genera from 2003-2023. *Vipera* (black), *Montivipera* (red), *Macrovipera* (blue) and *Daboia* (orange) from different geographical areas within 2003 to 2023. The bottom map shows the zoomed detailed overview of venomics studies on Turkish viper taxa with the original studies. Investigated taxa in this study are shown by images of the corresponding snake. Samples/specimen of non-reported venom origin were allocated to the respective capital city of the country. Closely located samples were summed to disks of increasing size. All images by Bayram Göçmen, except *Daboia* by Mert Karış.

It is important not to generalize the venom composition of a single population to be *per se* representative for a whole species or subspecies. They should rather be considered as potential source for differing compositions, as it has been already reported for several vipers ^16,29,32,33^. Remarkably, a large number of species and most subspecies have never been analysed by state of the art approaches, like modern venomics, defined as the application of high-throughput methodologies to the analysis of an organism’s full venom arsenal ^16,34^. Investigating these neglected taxa will help to predict the effect of a snakebite envenoming, to optimize treatment strategies, but also unveil venom evolutionary ecology and guide biodiscovery ^29,35–39^. This can be achieved either through the isolation characterization and assessment of single toxins *in vitro* and *in vivo*, or by detailed and data intensive omics approaches, like genomics, transcriptomics and proteomics ^38,40–42^. Especially the proteomic bottom-up (BU) ‘snake venomics’ approach, a three-step protocol with a final HPLC (high performance liquid chromatography) linked high resolution mass spectrometry (HR-MS) peptide detection, gives insights into compositions and allows cross-study comparison ^43–45^. Therefore, it has been used to correlate snake venoms in larger biogeographic contexts ^16,46–48^.

On the border between Europe and Asia, Türkiye represents a hotspot of snake diversity, hosting members of all four Palearctic viper genera ^49^. The rich herpetofauna has more than ten venomous snake species, several of which with an unresolved taxonomic status ^18,50,51^. Similar to tropical and subtropical regions, snakebite represents a major health burden in Türkiye, but the exact magnitude remains unclear due to the lack of comprehensive data ^52–54^. Several studies address concrete numbers about snakebite envenoming in Türkiye, like Oto and Haspolat (2021) showed that in southeast regions alone 108 children has been hospitalized between 2006 and 2011, not including dry bites ^55^. In earlier studies Karakus *et al*. (2015) reported for Hatay, a single province in South Türkiye, 125 cases from 2006 to 2010 in total, while Cesaretli and Ozkan (2010) listed 550 recorded envenomation by the National Poison Information Center (NPIC) across the whole country from 1995 to 2004 ^52,56^.While awareness of snakebite grows and the still underestimated numbers show the danger of envenomation, the species responsible for a bite are often not known. It is therefore necessary to investigate the range of venomous snakes in the country and the extent to which their venoms are composed. In the last decade, a few of these Turkish species have been studied using modern venomics approaches (**Figure 1**). These include representatives of Viperinae (*Vipera*, *Montivipera* and *Macrovipera*), as well as Morgan’s desert cobra, *Walterinnesia morgani* as the only elapid within this region ^16,57–63^. In this light, it is unfortunate that especially the Turkish taxa of highest medical significance remains virtually unstudied. Therefore, the venom composition and the potentially unfolding effects of envenoming stemming from such components are largely unknown hindering therapeutically care of snakebite victims.

Here, we set out to fill this knowledge gap and investigate the venom composition of seven Turkish viper taxa, many of which being recognized as threats to health. Specifically, we investigate representatives of each Turkish viperine genus by a combination of BU snake venomics and top-down (TD) proteomics including peptidomics ^64,65^. We describe for the first time the venom composition of the Baran’s adder *Vipera berus barani* (Böhme & Joger, 1983), an endemic subspecies of the adder located on the north of Türkiye, and the Darevsky’s viper *Vipera darevskii* (Vedmederja *et al.*, 1986), a small critically endangered viper living in close proximity to the Turkish-Georgian-Armenian border ^66,67^. Furthermore, aiming to gain a deeper understanding of the mountain viper venoms, we provide insights into three closely related and highly dangerous *Montivipera xanthina* complex: *Montivipera bulgardaghica bulgardaghica* (Nilson & Andren, 1985) and *Montivipera bulgardaghica albizona* (Nilson *et al*., 1990), as well as the Ottoman Viper *M. xanthina* (Gray, 1849) ^49,50,68–70^. The other two medical relevant genera are represented by one blunt-nosed viper subspecies *Macrovipera lebetinus obtusa* (Dwigubsky, 1832) and the Palestine viper *Daboia palaestinae* (Werner, 1938) ^71,72^. Our sample derived from the most northern, newly described Anatolian specimen of *D. palaestinae*, which venom of this region has been unknown until now ^73^.

By extensive modern venomics analysis we double the number of reported Turkish vipers venom compositions and gain novel insights in the venom variation of the four Old World viper genera *Vipera*, *Montivipera*, *Macrovipera* and *Daboia* on the proteomics as well as peptidomics level.

## 2. MATERIALS AND METHODS

### 2.1. Origin of snake venoms

All snakes were wild caught within Türkiye, the collections were approved with ethical permissions (Ege University, Animal Experiments Ethics Committee, 2010-2015) and special permissions (2011-2015) for field studies from the Republic of Türkiye, Ministry of Forestry and Water Affairs were received. For a detailed list of permission numbers, locations of collection and further venom pool information, see **Supplementary Table S1**.

### 2.2. Bottom-up proteomics - Snake Venomics

#### 2.2.1. Venom fractionation

For the analysis of each venom pool, 1 mg of lyophilized venom was dissolved to a final concentration of 10 mg/mL in aqueous 5% (*v*/*v*) acetonitrile (ACN) with 1% (*v*/*v*) formic acid (HFo) and centrifuged for 5 min at 10,000 × g. The supernatants were fractionated on a reversed-phase Supelco Discovery BIO wide Pore C18-3 (4.6 × 150 mm, 3 µm particle size) column operated by a HPLC Agilent 1200 (Agilent Technologies, Waldbronn, Germany) chromatography system. The following gradient with ultrapure water with 0.1% (v/v) HFo (solvent A) and ACN with 0.1% (v/v) HFo (solvent B) was used at 1 mL/min, with a linear gradient between the time points, given at min (B%): 0–5 (5% const.), 5–100 (5 to 40%), 100–120 (40 to 70%), 120–130 (70% const.), and 5 min re-equilibration at 5% B. The chromatography runs were observed by a diode array detector (DAD) at λ = 214 nm detection wavelength. Samples were collected through time-based fractionation (1 fraction/min) and combined peak fractions were dried in a centrifugal vacuum evaporator.

Peaks later than 25 min were further processed by the snake venomics steps of gel separation and tryptic digest, peaks with earlier retention times (R_t_) are known for their low molecular mass peptide content and were directly sent to the LC-MS. The viperine abundant tripeptide pEKW (with pE for pyroglutamate) signal at around 25 min was set as benchmark.

#### 2.2.2. SDS-PAGE profiling and tryptic digestion

The dried venom fractions were redissolved in 10 µL reducing 2× sodium dodecyl sulfate (SDS) sample buffer (125 mM Tris HCl pH 6.8, 4% (w/v) SDS, 17.5% (w/v) glycerol, 0.02% (w/v) Bromphenol blue and 200 mM freshly prepared dithiothreitol DTT in ultra-pure (MQ) water), heated for 10 min at 95 °C, fully loaded and separated using a 12% SDS-PAGE (SurePage Bis-Tris, Genscript, Piscataway, NJ, USA) run with MES buffer (50 mM 2-(*N*-morpholino)ethane sulfonic acid (MES), 50 mM Tris base, 1 mM EDTA, 0.1% (w/v) SDS, stored in brown glass flasks at 4°C) at 200 V for 21 min. A PageRuler Unstained Protein Ladder (Thermo Scientific, Waltham, MA, USA) was used as protein mass standard. Gels were three times short-washed with water. Proteins were fixed with preheated (50-60 °C) fixation buffer three times for 10 min each (aqueous, 40% (v/v) methanol, 10% (v/v) acetic acid), stained for 45 min in preheated (50-60 °C) fast staining buffer (aqueous, 0.3% (v/v) HCl 37%, 100 mg/L Coomassie 250G) under constant mild shaking, and kept overnight at 4 °C in storage buffer (aqueous, 20% (v/v) methanol, 10% (v/v) acetic acid) for destaining. The cleaned gels were then scanned for documentation and quantification. Gel pieces with single protein bands were cut, dried with 500 µL ACN, and stored at −20 °C without ACN until tryptic digestion. The disulfide bridges were reduced with 30 µL freshly prepared DTT (100 mM in 100 mM ammonium hydrogen carbonate (ABC) per gel band) for 30 min at 56 °C and dried with 500 µL ACN for 10 min before removing the supernatant. Cysteines were alkylated with freshly prepared iodoacetamide (55 mM in 100 mM ABC) for 20 min at room temperature in the dark to protect the reduced thiols from oxidation and washed with 500 µL ACN for 2 min. before removing the supernatant. Gel samples were dried again with 500 µL ACN for 15 min, ACN removed, followed by 30 min incubation on ice with 30 µL freshly activated trypsin (13.3 ng/µL, 10% (v/v) ACN in 10 mM ABC; Pierce trypsin, Thermo, Rockfeld, IL, USA). When necessary, additional volumes of trypsin were added, so that the gel piece was still covered in buffer. All samples were incubated for 90 min on ice, 20 µL ABC buffer (10 mM) was added, and were incubated overnight at 37 °C. Peptides were extracted with 100 µL pre-warmed elution buffer (aqueous, 30% (v/v) ACN MS grade, 5% (v/v) HFo) at 37 °C for 30 min. The supernatant was transferred into a separate microtube, vacuum-dried and if possible directly prepared for the LC-MS/MS measurement, else samples were stored at −20 °C.

#### 2.2.3. Mass spectrometry

For the MS analysis, the extracted and dried tryptic peptides were re-dissolved in 30 µL aqueous 3% (*v*/*v*) ACN with 1% (*v*/*v*) HFo, and 20 µL of each was injected into an LTQ Orbitrap XL mass spectrometer (Thermo, Bremen, Germany) via an Agilent 1260 HPLC system (Agilent Technologies, Waldbronn, Germany) using a reversed-phase Grace Vydac 218MS C18 (2.1 × 150 mm; 5 μm particle size) column. The following gradient with ultrapure water with 0.1% (*v*/*v*) HFo (solvent A) and ACN with 0.1% (*v*/*v*) HFo (solvent B) was used at 0.3 mL/min, with a linear gradient between the time points, given at min (B%): 0–1 (5% const.), 1–11 (5 to 40%), 11–12 (40 to 99%), 12–13 (99% const.), and 2 min re-equilibration at 5% B. The parameters in the ESI positive modus were as follows: 270 °C capillary temperature, 45 L/min sheath gas, 10 L/min auxiliary gas, 4.0 kV source voltage, 100.0 µA source current, 20 V capillary voltage, 130 V tube lens. FTMS measurements were performed with 1 μ scans and 1000 ms maximal fill time. AGC targets were set to 10^6^ for full scans and to 3 × 10^5^ for MS2 scans. MS2 scans were performed with a mass resolution (R) of 60,000 (at *m*/*z* 400) for *m*/*z* 250–2000. MS2 spectra were obtained in data-dependent acquisition (DDA) mode as top2 with 35 V normalized CID energy, and 500 as the minimal signal required with an isolation width of 3.0. The default charge state was set to z = 2, and the activation time to 30 ms. Unassigned charge states and charge state 1 were rejected for tryptic digest peptides, for direct submitted fractions from the initial HPLC run all charge states were measured.

### 2.3. Bottom-up data analysis

The BU LC-MS/MS data RAW files were converted into the MASCOT generic file (MGF) format using MSConvert (version 3.0.10577 64-bit) with peak picking (vendor msLevel = 1−) ^74^. For an automated database comparison, files were analysed using pFind Studio ^75^, with pFind (version 3.1.5) and the integrated pBuild, with the following parameters: MS Data (format: MGF; MS instrument: CID-FTMS); identification with Database search (enzyme: Trypsin KR_C, full specific up to 3 missed cleavages; precursor tolerance +20 ppm; fragment tolerance +20 ppm); open search setup with fixed carbamidomethyl [C] and Result Filter (show spectra with FDR ≤ 1%, peptide mass 500–10,000 Da, peptide length 5–100 amino acids, and show proteins with number of peptides >1 and FDR ≤ 1%). The used databases included UniProt ‘Serpentes’ (ID 8750, reviewed, canonical and isoform, 2640 entries, last accessed on 8^th^ April 2021 via https://www.uniprot.org/) and the Common Repository of Adventitious Proteins (215 entries, last accessed on 10 February 2022; available at https://www.thegpm.org/crap/index.html). The results were batch-exported as PSM score of all peptides identified with pBuild and manually cleared from decoy entries, contaminations, and artifacts to generate the final list of unique peptide sequences per sample with the best final score. For a second confirmation of identified sequences, all unique entries were analysed using BLAST search with blastp against the non-redundant protein sequences (nr) of the “Serpentes” (taxid: 8570) database ^76,77^. In case of non-automatically annotated band identity, files were manually checked using Thermo Xcalibur Qual Browser (version 2.2 SP1.4), *de novo* annotated, and/or compared on MS1 and MS2 levels with other bands to confirm band and peptide identities. Deconvolution of isotopically resolved spectra was carried out by using the XTRACT algorithm of Thermo Xcalibur.

### 2.4. Top-down proteomics

For the denaturing TD analysis, 100 µg of lyophilized venom was dissolved to a final concentration of 10 mg/mL in aqueous 1% (*v*/*v*) HFo and centrifuged for 5 min at 20,000 × g. The supernatant was mixed with 30 μL of citrate buffer (0.1 M, pH 3.0) and split into two aliquots. The first aliquot was mixed 10 μL of 0.5 M tris(2-carboxyethyl)phosphine (TCEP), for reduction of disulfide bonds, and incubated for 30 min at 65 °C. The second was supplemented with 10 μL of ultrapure water and will be referred as non-reduced sample. Both samples were centrifuged for 5 min at 20,000 × g and 10 µL of each was injected into an Q Exactive HF mass spectrometer (Thermo, Bremen, Germany) via a Vanquish ultra-high performance liquid chromatography (UHPLC) system (Agilent Technologies, Waldbronn, Germany) using a reversed-phase Supelco Discovery BIO wide C18 (2.0 × 150 mm; 3 μm particle size; 300 Å pore size) column thermostated at 30 °C. The following gradient with ultrapure water with 0.1% (*v*/*v*) HFo (solvent A) and ACN with 0.1% (*v*/*v*) HFo (solvent B) was used at 0.4 mL/min, with a linear gradient between the time points, given at min (B%): 0–6 (5% const.), 6–25 (5 to 40%), 25–30 (40 to 70%), 30–35 (70% const.), and 5 min re-equilibration at 5% B. The parameters in the ESI positive modus were as follows: 265.50 °C capillary temperature, 50.00 AU sheath gas, 12.50 L/min auxiliary gas, 3.50 kV source voltage, 100.00 µA source current. FTMS measurements were performed with 1 μ scans and 1000 ms maximal fill time. MS2 scans were performed with a mass resolution (R) of 140,000 (at *m*/*z* 200). MS2 spectra were obtained in DDA mode as top3 with 30% normalized high energy C-trap dissociation (HCD) and an isolation window of *m/z* 3.0. The default charge state was set to z = 6, and the activation time to 30 ms. Unassigned charge states and isotope states were rejected for MS2 measurements.

### 2.5 Top-down data analysis

The TD LC-MS/MS Thermo RAW data were converted to a centroided mass spectrometry data format (mzML) using MSConvert (version 3.0.10577 64-bit) with peak picking (vendor msLevel = 1−) and further analyses by TopPIC ^74,78^. The mzML data were deconvoluted to a MSALIGN file using TopFD (http://proteomics.informatics.iupui.edu/software/toppic/; version 1.6.5) with a maximum charge of 30, a maximum mass of 70 000 Da, an MS1 S/N ratio of 3.0, an MS2 S/N ratio of 1.0, an *m*/*z* precursor window of 3.0, an *m*/*z* error of 0.02 and HCD as fragmentation ^79^. The final sequence annotation was performed with TopPIC (http://proteomics.informatics.iupui.edu/software/toppic/; version 1.6.5) with a decoy database, maximal variable PTM number 3, 10 ppm mass error tolerance, 0.01 FDR cutoff, 1.2 Da PrSM cluster error tolerance, and a maximum of 1 mass shifts (±500 Da), and a combined output file for the non-reduced and reduced samples of a venom pool ^78^. Spectra were matched against the UniProt ‘Serpentes’ database (ID 8750, reviewed, canonical and isoform, 2749 entries, last accessed on 11^th^ October 2023 via https://www.uniprot.org/), manually validated, and visualized using the MS and MS/MS spectra using Qual Browser (Thermo Xcalibur 2.2 SP1.48). The XTRACT algorithm of Thermo Xcalibur was used to deconvolute isotopically resolved spectra.

### 2.6. Intact mass profiling and peptidomics

The TD RAW data were manually screened in the Qual Browser (Thermo Xcalibur 2.2 SP1.48) for an overview of abundant intact protein and peptide masses. They were correlated to the previous peak annotation and identification by snake venomics as well as used for the counting of disulfide bridges between the non-reduced and reduced TD RAW samples. Spectra of multiple charges were isotopically deconvoluted by using the XTRACT algorithm of Thermo Xcalibur. Masses in this study are given in the deconvoluted average *m/z* (with z=1), if not stated otherwise. Monoisotopic masses are also given with z=1. In case of abundant non-TD-annotated peptides, masses were manually checked using Thermo Xcalibur Qual Browser (version 2.2 SP1.4), the peptide sequences were manually *de novo* annotated by the MS/MS spectra and the *m/z* peaks cross-confirmed by in silico fragmentation using MS-Product of the ProteinProspector (http://prospector.ucsf.edu, version 6.4.9) ^80^.

### 2.7. Proteome quantification

The used quantification protocol is based and adapted to the common three-step ‘snake venomics’ approach as summarised in Calvete *et al.* 2023 ^81^. The comparable approach determine a toxin family abundance in the venom as the sum of all its normalized toxin abundances *T*:

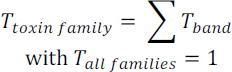

The normalized toxin abundance within a single protein band *T_band_* is calculated with the normalized values of the RP-HPLC peak integral *P* measured at 214 nm, the densitometric gel band intensity *D* and if necessary the relative MS ion intensity *M* of the most abundant and identified peptides:

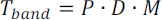

For the peak quantification after blank run subtraction, the HPLC separation chromatogram fractions were integrated as area under the curve *P_peak_* in ratio to the total sum of all peaks:

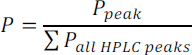

For the densitometric quantification of a single SDS band, the non-highly compressed gel scan (here in PNG format) was processed by Fiji ^82^. The colour depth was set to 8bit grayscale and inverted to integrate former darker bands with higher values. The band area *A_band_* and the corresponding integrated band densities *D_band_* were measured for each band, as well as a corresponding background areas *A_bg_* and integrated band densities *D_bg_*. By removing the proportion of background, we calculated the normalized gel band intensity *D* for each toxin band in the gel:

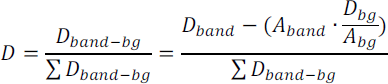

In case of multiple toxin identification within a single band, single normalized toxin abundances *M* were estimated based on the ion intensity sum of the three most intensive peptide ions of one toxin from *M_3_* in relation to the sum of all top3 toxin ions from the other co-migrated toxins families within this MS sample, as summarised in Calvete *et al.* 2023 ^81^:

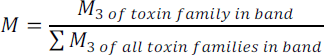

In total, band identification based on the BU, TD and peptidomics results, in comparison to the IMP and the apparent masses of the SDS bands.

### 2.8. Online proteome search

To identify relevant publications for the comparison of venom compositions the review of Damm *et al*. (2021) was used as template and database for Old World vipers (Squamata: Serpentes: Viperidae: Viperinae) venoms ^16^. We used the identical selection criteria parameters with two modifications. Firstly, the genera, species, and subspecies taxa search were limited to Palearctic vipers of the genus *Vipera*, *Montivipera*, *Macrovipera* and *Daboia,* and the investigated time window was continued from 1^st^ January 2021 until 31^st^ December 2023. Therefore the PubMed database (https://pubmed.ncbi.nlm.nih.gov/) of the National Centre of Biotechnology Information (NCBI), Google (https://www.google.com/) as well as Google Scholar (https://scholar.google.com/) has been searched as described earlier and the results were screened manually for proteomic studies ^16^.

### 2.9. Data accessibility

Mass spectrometry proteomics data have been deposited in the deposited to the PRIDE Archive (http://www.ebi.ac.uk/pride/archive/) via the PRIDE partner repository with the data set identifier Mass spectrometry proteomics data have been deposited with the ProteomeXchange Consortium10 via the MassIVE partner repository (https://massive.ucsd.edu/) under the bottom-up and top-down project names “Snake venom proteomics of seven taxa of the genera *Vipera*, *Montivipera*, *Macrovipera* and *Daboia* across Turkiye/Turkey” with the dataset identifiers “MSV000094228” and “MSV000094229”, respectively, as well as in the Zenodo repository (https://zenodo.org) under the project name “DATASET - Mass Spectrometry - Snake venom proteomics of seven taxa of the genera *Vipera*, *Montivipera*, *Macrovipera* and *Daboia* across Türkiye” with the dataset identifier 10.5281/zenodo.10683187 ^83^.

## 3. RESULTS

The venom proteomes of seven Palearctic viper taxa of Turkish origin were profiled by the snake venomics approach (**Figure 2,3 and 5, Supplementary Figures S1-S7**). For a comprehensive analysis each venom was additionally investigated by non-reduced and reduced top-down MS, including intact mass profiling and peptidomics. All identified toxins and homologs are in detail listed in the supplements (**Supplementary Tables S3-S9**). Four venom proteomes represent first descriptions for these snake taxa (*V. b. barani*, *V. darevskii*, *M. b. albizona* and *M. xanthina*), two have never been investigated before by extensive snake venomics for Turkish populations (*M. l. obtusa* and *D. palae*stinae) and one is an in-depth reanalysis in order to identify >20% of unknown proteins from a previous study (*M. b. bulgardaghica*, identical pool) ^58^. In general, the seven proteomes largely conform to the previously proposed compositional family trends of toxins in viperine venoms ^16^. Accordingly, viperine venoms can be categorized into typical major-, secondary-, and minor toxin families. For those, the following abundance ranges were identified for the herein analyzed venoms:

- major toxin families: snake venom metalloproteinases (svMP, <1-34%) including disintegrin-like/cysteine-rich (DC) proteins; snake venom phospholipases A_2_ (PLA_2_, 8-18%); snake venom serine proteases (svSP, 10-46%); C-type lectin-related proteins and snake venom C-type lectins (summarized as CTL, 3-20%),
- secondary toxin families: disintegrins (DI, 0-15%); L-amino acid oxidases (LAAO, 2-4%); cysteine-rich secretory proteins (CRISP, 0-13%), vascular endothelial growth factors F (VEGF, 0-12%), Kunitz-type inhibitors (KUN, 0-9%),
- minor toxin families: i.a. 5′-nucleotidases (5N, 0.1-0.8%); nerve growth factors (NGF, 0.3%); phosphodiesterases (PDE, 0.2%).

Members of rare families in Viperinae venoms, like glutaminyl cyclotransferases (EC 2.3.2.5) or aminopeptidases (EC 3.4.11.-), have not been detected in the herein studied venoms. In the following section, each snake venom composition will be described and the proteomes will be discussed on a genus-wide comparison. Furthermore, a variety of peptides (9-19%) have been observed in the venoms and will be highlighted later in detail separately.

### 3.1. Vipera berus barani *and* V. darevskii

**Figure 2.**
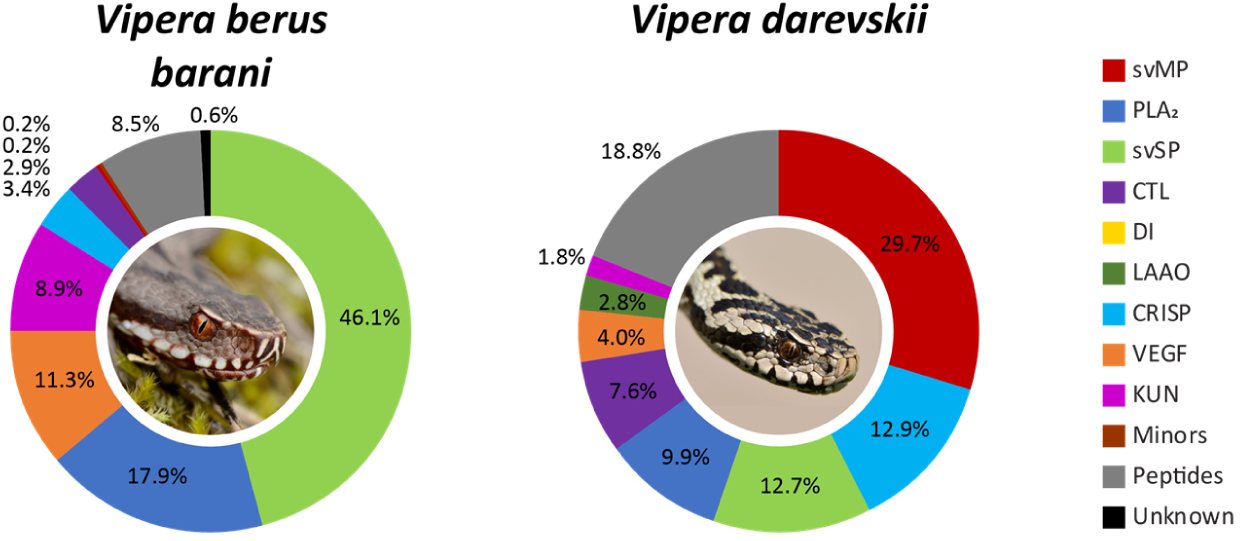
*Vipera* venom compositions of *V. b. barani* and *V. darevskii*. The venom proteomes of two *Vipera* taxa from Türkiye have been quantified by the combined snake venomics approach via HPLC (λ = 214 nm), SDS (densitometry) and MS ion intensity, including TD proteomics. Toxin families are arranged clockwise by abundances, followed by peptides (grey) and non-annotated parts of the venom (unknown, black). Images by Bayram Göçmen.

With *V. b. barani* and *V. darevskii* two different taxa of the *Vipera* subclade *Pelias* have been analysed in this study (**Figure 2, Supplementary Table S3/4, S10/11, S17/18**). The *V. b. barani* crude venom HPLC profile lacks abundant peaks at R_t_ >90 min, corresponding to a higher ACN gradient (**Supplementary Figure S1**). In viperines, those peaks include normally P-III svMP and have been observed in all other venoms examined within this study. In contrast, *V. b. barani* lacks those late-eluting peaks, and svMP are surprisingly underrepresented and correspond to only 0.2% of the venom. They were identified as members of the P-III subfamily and accordingly no DI were observed.

On the other side, the venom profile has a complex peak structure in the chromatogram between 75 to 90 min (F27-38) and svSP were identified as the most abundant toxin family. The fractions (F) F27-45 contain svSP of up to 32 kDa and the IMP revealed *m/z* 30327.40 and *m/z* 30909.67 as the most abundant average svSP masses. Both masses appeared in groups of peaks, based on the variable *N*-glycosylation. The clearest signals had mass shifts of Δ203 Da and Δ406 Da, indicating at least two *N*-acetylhexosamines (HexNAc, 203.08 Da) in the glycosylation tree. By BU, nikobin was identified as homolog in most of the fractions with the highest peptide coverage. The remaining svSP were identified as homologs to the hemotoxic factor V-activating enzyme (RVV-V, *D. siamensis*) or svSP homolog 2 (*M. lebetinus*).

A combination of basic, neutral and acidic PLA_2_ (18%) formed the second most abundant toxin family and all PLA_2_ in the *V. b. barani* venom were identified as neurotoxic homologs, like ammodytin L and ammodytoxin C via BU proteomics ^84,85^. By TD proteomics proteoforms of ammodytin (*m/z* 13553.88, 13676.39, 13692.84) and ammodytoxin (*m/z* 13742.19, 13773.18, 13856.25) were annotated and the PLA_2_ conserved seven intramolecular disulfide bridges could be confirmed between the reduced and non-reduced samples (**Supplementary Table S17**). The following most abundant toxin families were VEGF (11%), mostly vammin-1’ related, and KUN (9%) formed by a single serine protease inhibitor ki-VN (*m/z* 7594.47) with three TD confirmed disulfide bridges. Further toxin families are CRISP (3%), with a single dominant band in F24/25 of *m/z* 24599.42, CTL (3%), PDE (0.2%) and LAAO in small traces (44c; ∼55 kDa). Abundant peptides signals of low molecular sizes have been identified by MS2 as pERRPPEIPP (*m/z* 1072.59) and pERWPGPKVPP (*m/z* 1144.62), beside two tripeptidic svMP inhibitors (svMP-i) pEKW (*m/z* 444.22) and pERW (*m/z* 472.23).

The second *Vipera* venom investigated in this study stems from *V. darevskii*. It largely follows the classical Viperinae composition and is characterized by a high abundances of svMP (30%, P-III svMP only), PLA_2_ (10%), svSP (13%) and CTL (8%) as major toxin families.

The main PLA_2_ are acidic homologs to the toxins from *V. ammodytes* and *V. renardi*, such as myotoxic ammodytin l1, as well as MVL-PLA2 and VpaPLA2 from *Daboia* and *Macrovipera* species. One third of the svSP (4% of the total venom) shared the highest similarities with anticoagulant active homologs of *V. ammodytes* and *M. lebetinus*, while the remaining 9%, all eluting >80 min, were matched to sequences from *V. berus* (nikobin) and *V. anatolica senliki*. The CRISP (13%) toxins are second most abundant and interestingly a strong signal for a CRISP fragment has been observed with a monoisotopic mass of *m/z* 6414.61 eluting at 11 min in the non-reduced, non-digested venom. Its reduced monoisotopic signal of *m/z* 6424.68 could be annotated by TD as the *C*-terminal fragment of CRVP_VIPBN, a CRISP from *V. berus nikolskii*, with a single oxidation (+15.99 Da). The mass shift of Δ10.065 Da indicates five disulfide bridges through ten Cys in the sequence. Several further secondary toxin families were identified, like VEGF (4%), LAAO (3%) and KUN (2%), but no DI nor any minors or rare were detected. The peptides (19%) are dominated by a single svMP-i (pEKW) fraction with over 11% of the whole venom proteome of *V. darevskii*. Furthermore, 3% could be assigned to the de novo annotated peptide pENWPGPK (*m/z* 809.39).

### 3.2. Montivipera bulgardaghica *ssp. and* M. xanthina

**Figure 3.**
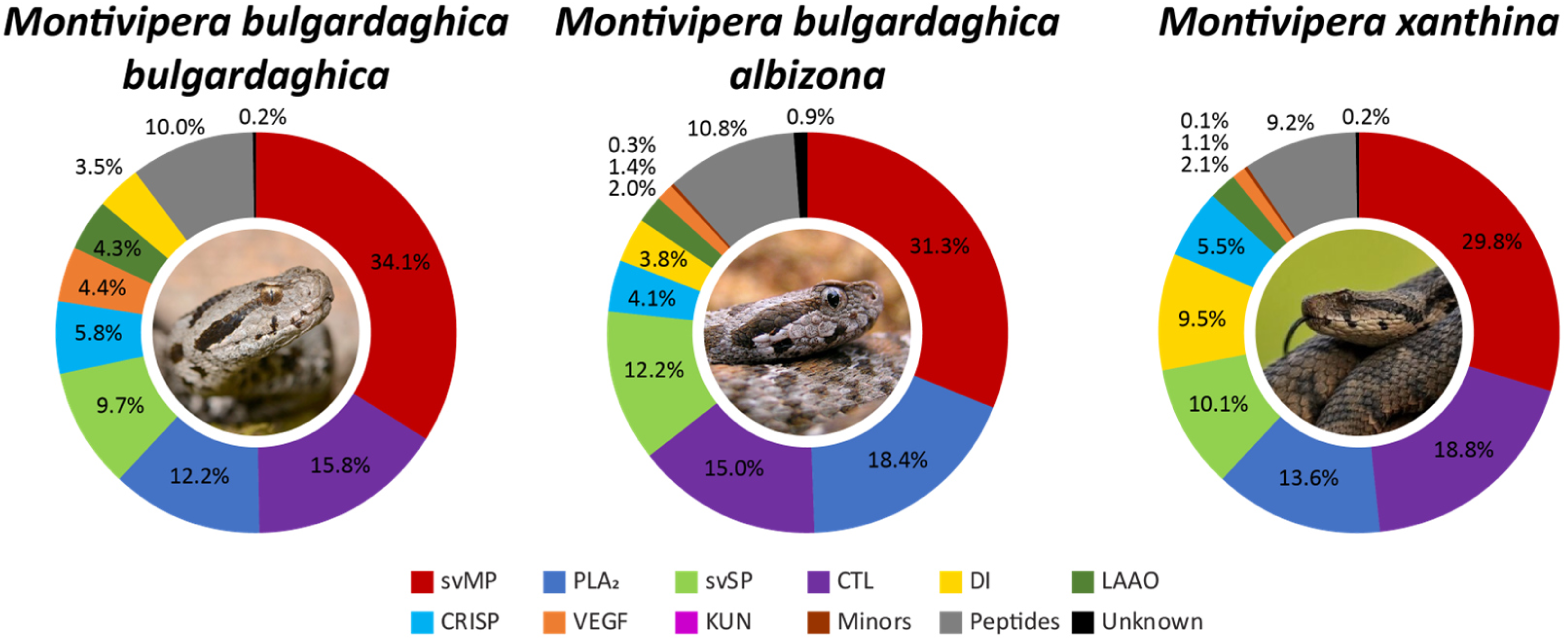
*Montivipera* venom compositions of *M. b. bulgardaghica*, *M. b. albizona* and *M. xanthina*. The venom proteomes of three *Montivipera* taxa from Türkiye have been quantified by the combined snake venomics approach via HPLC (λ = 214 nm), SDS (densitometry) and MS ion intensity, including TD proteomics. Toxin families are arranged clockwise by abundances, followed by peptides (grey) and non-annotated parts of the venom (unknown, black). Images by Bayram Göçmen.

The genus of *Montivipera* is represented in this study by three different taxa, two *M. bulgardaghica* subspecies (*M. b. bulgardaghica*, *M. b. albizona*) and *M. xanthina* (**Figure 3, Supplementary Table S5-7, S12-15, S19-21**). A chromatogram comparison revealed 53 signal groups in the venom of *M. b. bulgardaghica*, 50 for *M. b. albizona* and 42 for *M. xanthina* (**Supplementary Figures S3-S5**) The profiles between the *M. bulgardaghica* ssp. had higher similarities in the chromatograms of the first 75 min compared to *M. xanthina*, while eluting profiles between 80 to 110 min of all three venoms had exhibited striking similarities (**Figure 4**).

**Figure 4.**
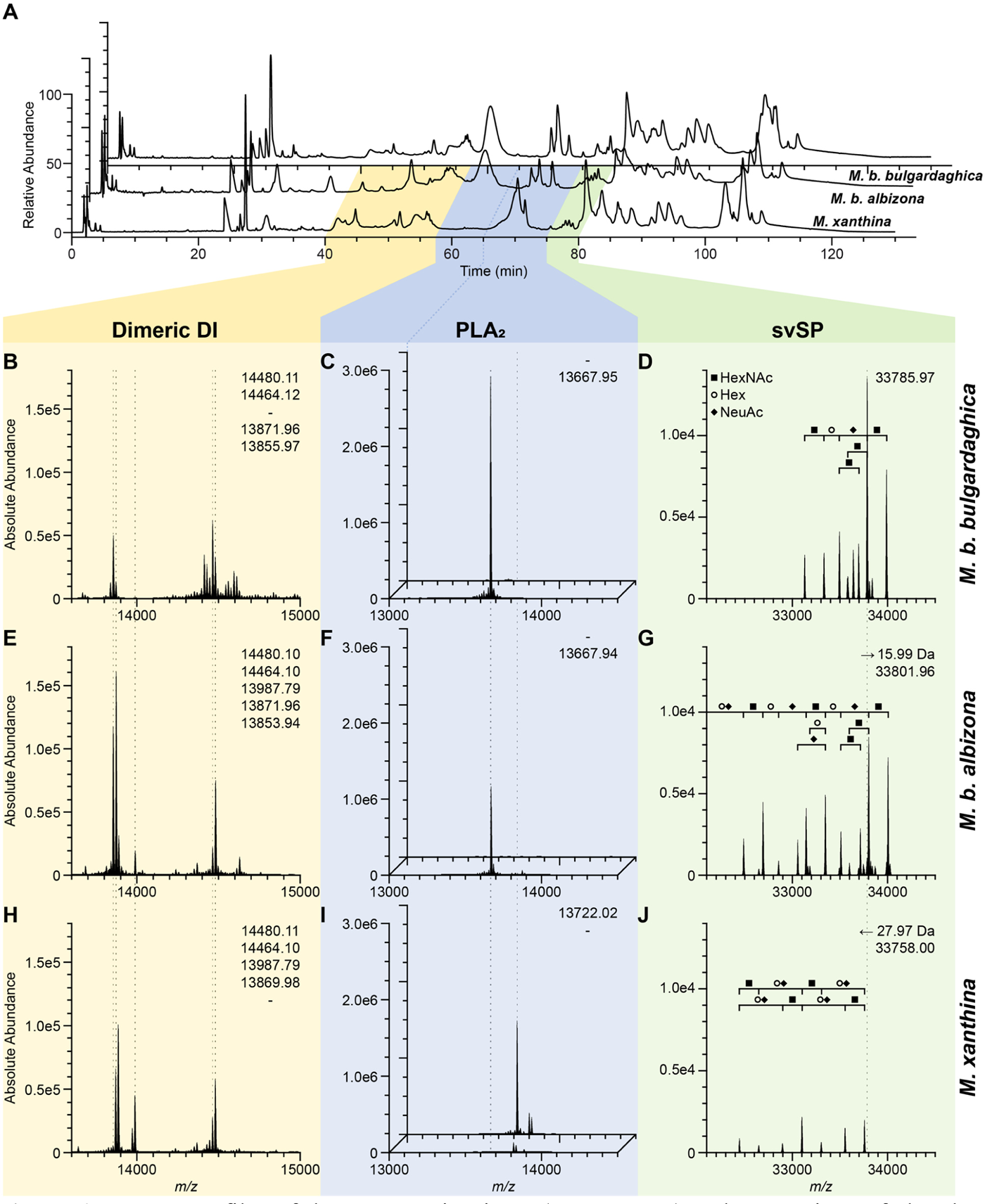
Venom profiles of three mountain vipers (*Montivipera*) and comparison of abundant toxins. (**A**) Chromatogram of the venoms from *M. b. bulgardaghica* (top/back; **B-D**), *M. b. albizona* (middle; **E-G**) and *M. xanthina* (bottom/front; **H-J**) with λ = 214 nm. (**B-J**) Exemplary main toxin families were investigated by non-reduced intact mass profiling (IMP) at their corresponding top-down proteomics retention times set in correlation to the snake venomics HPLC profile. The deconvoluted main toxin masses (dashed lines) are compared for five dimeric DI (**B, E, H** at 11.4-15.2 min IMP RT) and two PLA_2_ at two different times (**C,F,I** at front 15.3-18.0 min and back 18.0-19.7 min IMP RT). Begin of the second PLA_2_ time windows in (**A**) is connected (dark blue line) the corresponding IMP (back of **C,F,I**). A svSP (**D,G,J** at 20.5-21.2 min IMP RT) shows small mass shifts but similar glycosylation components: HexNAc (*N*-acetylhexosamines, filled square), Hex (hexose, circle), NeuAc (*N*-acetyl neuraminic acid, filled rhombus). Abbreviations: DI, disintegrins (yellow); PLA_2_, phospholipase A_2_ (blue); svSP, snake venom serine protease (green).

The semi-quantification by snake venomics shows high similarities in the major toxin abundances. In all three *Montivipera* venoms different svMP (30-34%) dominate, mostly P-III svMP to a smaller extend of DC proteins (2-4%), followed by CTL (15-19%) (**Figure 3**). Each venom had three main fractions collected between 82-104 min with abundant CTL bands in the reduced SDS PAGE at 12 to 15 kDa, each. This is consistent with their multimeric structure ^86^. The observed tryptic peptides sequences were homolog to *M. lebetinus* toxins in all three snakes: Snaclec A11/A1/B9 (82 min), Snaclec A16/B7/B8 (88 min) and C-type lectin-like protein 3A (104 min). At 104 min also Snaclec 3 from *D. siamensis* has been identified and the TD annotation confirmed the presence of Snaclec A14 homologs (*M. lebetinus*) in each of the *Montivipera* venoms.

The PLA_2_ (12-18%) differ between the species. The acidic phospholipase A_2_ Drk-a1 homolog, from *D. russelii*, is the main representative in both, *M. b. bulgardaghica* (11%) and *M. b. albizona* (12%), with *m/z* 13667.95 and *m/z* 13667.94, respectively (**Figure 4C,F,I**). The PLA_2_ were detected in a single dominant peak at R_t_ = 62 min, at which the *M. xanthina* chromatogram had only a flat broad signal (F22). In the *M. xanthina* composition this fraction has been identified by BU as a coelution of NGF (0.1%) and PLA_2_ (1.3%) with a mass of *m/z* 13833.24. Its main PLA_2_ eluted a few minutes later at ∼70 min forming a strong signal (F23-25), which in turn was absent in the first two profiles. In *M. xanthina* a different main acidic PLA_2_ homolog with *m/z* 13722.02 has been observed. It represents over 8% of the whole venom (**Figure 4C,F,I**). The remaining 3% were formed by basic PLA_2_, which were only be detected in traces within the two *M. bulgardaghica* subspecies.

Within all three HPLC profiles a group of close eluting peaks has been detected at <80 min, which is typical for svSP in viper venoms bearing an extensive glycosylation. BU proteomics confirmed the presence of svSP and the IMP revealed several molecular masses around 33 kDa. The main svSP masses differ within the genus of *Montivipera*, but all peaks are closely related with mass shifts of Δ15.99 Da (O) between *M. b. bulgardaghica* (*m/z* 33785.97) and *M. b. albizona* (*m/z* 33801.96), and Δ27.97 Da (CO) between *M. b. bulgardaghica* and *M. xanthina* (*m/z* 33758.00) (**Figure 4D,G,J**). All three had peak patterns of same distances and revealed so similar consecutive glycosylations, with observed mass shifts of Δ203 Da (HexNAc, 203.08 Da), Δ162 Da (hexose Hex, 162.06 Da) and Δ291 Da (*N*-acetyl neuraminic acid NeuAc, 291.10 Da) (**Figure 4D,G,J**).

Secondary toxin families were identified at lower abundances: DI (4-10%), CRISP (4-6%), LAAO (2-4%) and VEGF (1-4%) of which all belong to the vammin/ICCP-type ^87^, but no KUN have been detected in any *Montivipera* venom. In total, eleven different abundant masses could be identified as heterodimeric DI around 14 kDa, and while monomeric DI of various lengths from 4 to 8 kDa are known to appear in viper venoms, none of these have been observed in the herein analyzed *Montivipera* venoms. *M. xanthina* showed with 9.5% more than twice the amount of DI than *M. b. bulgardaghica* (3.5%) and *M. b. albizona* (3.8%). Only two abundant dimeric DI are shared across all three venoms. They have molecular masses of *m/*z 14480.10 and *m/z* 14464.11 with Δ15.99 Da (O) (**Figure 4B,E,H**), and TD revealed the two subunits as homologs of the close related taxa: lebein and VLO5 (both *M. lebetinus*) and EMF10 (*Eristicophis macmahoni*). The sum of their corresponding exact masses gives proof of the characteristic ten disulfide bridges (twice four intra-plus two interchains). The other ten dimeric DI were either detected in two of the three vipers, or unique for one of them. For example, both *M*. *bulgardaghica* ssp. shared *m/z* 13871.96, while *m/z* 13987.79 has been only observed for *M. b. albizona* and *M. xanthina* (**Figure 4B,E,H**). The other dimeric masses between the three venoms show differences of a few Dalton, due to the high similarity of the monomeric subunit, similar to the later described *M. l. obtusa*.

The three CRISP containing peaks eluted contemporaneous in the *Montivipera* venoms at R_t_ = 70 min, with main representative masses of *m/z* 24816.46 in *M. b. bulgardaghica*, *m/z* 24806.44 in *M. b. albizona* and *m/z* 24666.54 in *M. xanthina*. The latter was also low abundant in *M. b. bulgardaghica* (*m/z* 24699.52), and not present in *M. b. albizona*. For the category of minor toxins only 5N (0.3%) were annotated by BU in the venom of *M. b. albizona* and NGF (0.1%) in *M. xanthina*.

The three herein analyzed *Montivipera* venoms contain a similar peptide part of around 10% and the svMP-i pEKW, pERW and pENW (*m/z* 430.17) could be identified in all of them as abundant components. The decapeptide pENWPSPKVPP (*m/z* 1132.55) and the two sequence related peptides pENWPSPK (*m/z* 839.41) and pENWPSP (*m/z* 711.31) were also prominent in each *Montivipera* peptidome as well as the glycine-rich peptide pEHPGGGGGGW (*m/z* 892.37).

### 3.3. Macrovipera lebetinus obtusa

**Figure 5.**
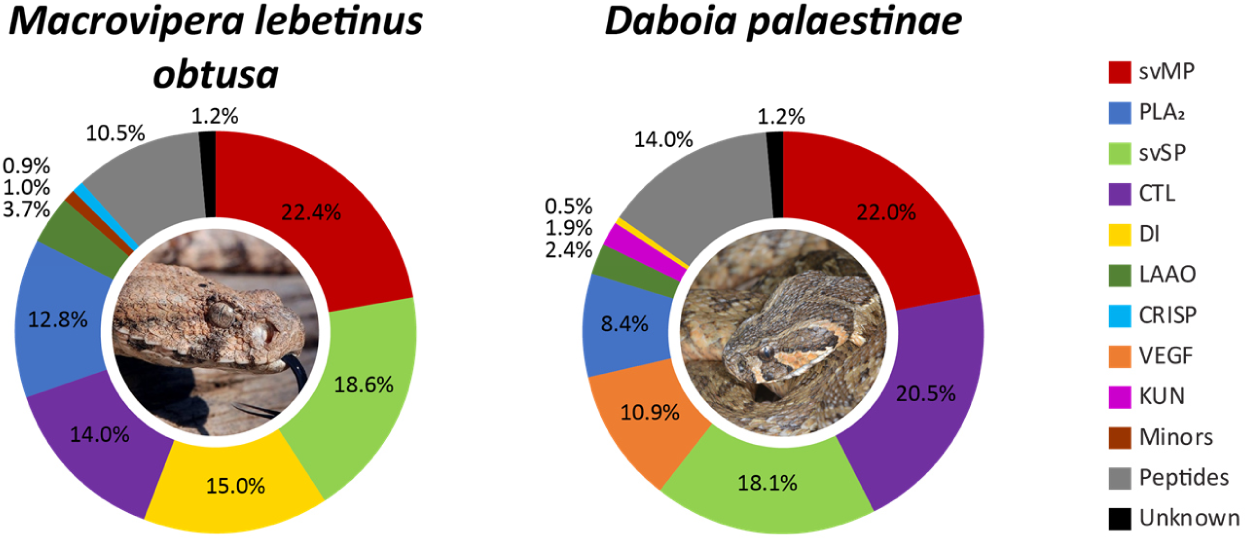
*Macrovipera* and *Daboia* venom compositions of *M. l. obtusa* and *D. palaestinae*. The venom proteomes of one *Macrovipera lebetinus* subspecies and one *Daboia* species from Türkiye have been quantified by the combined snake venomics approach via HPLC (λ = 214 nm), SDS (densitometry) and MS ion intensity, including TD proteomics. Toxin families are arranged clockwise by abundances, followed by peptides (grey) and non-annotated parts of the venom (unknown, black). Images by Bayram Göçmen (*Macrovipera*) and Mert Karış (*Daboia*).

The third Palearctic viper genus analysed was *Macrovipera*, also referred to as blunt-nosed vipers. Here, we examined the venom of *M. l. obtusa* (**Figure 5, Supplementary Table S8, S15, S22**). Its major toxins, including DI, forming 83% of the venom and are mostly composed of svMP (22%), with P-I (2%) and P-III svMP (12%). The DC proteins (8%), or P-IIIe svMP subfamily, account for >8% of the venom. They were identified as homologs of Leberagin-C (F22/23). The most abundant P-III svMP was the heavy chain of the coagulation factor X-activating enzyme VLFXA. It forms a heterotrimeric complex with the CTL light chains 1 and 2, annotated in F38 and F40. Further abundant svMP include the apoptosis inducing VLAIP-A/B (P-III) and lebetase (P-I). The svSP (19%) consist of different toxins, that has been previously described from the *Macrovipera* genus and a majority of the tryptic peptide sequences originated from the coagulant-active lebetina viper venom FV activator (VLFVA or LVV-V), followed by the α-fibrinogenase (VLAF), VLP2 and VLSP3. The third most common toxin family are DI (15%) and we could identify more than ten dimeric DI masses and several determined DI subunits within the Turkish *M. l. obtusa* venom (**Supplementary Table S24**). The main DI subunits, identified by TD and BU, are from known *Macrovipera* toxins, such as lebein-1, VB7A, VLO4, VLO5, VM2L2 or lebetase. This high variety of dimeric DI is also a result of mass shifts compared to known subunit sequences, originating from e.g. from oxidation (Δ15.99 Da), hydration (Δ18.01 Da), or the loss of terminal amino acids, e.g. seen *C*-terminal at lebein-1-alpha (-KD*^C^*) or *N*-terminal at VM2L2 (-*^N^*QNSGN) and VLO5B (-*^N^*M). It cannot be ruled out that some of these modifications are artifacts due to the analysis methods used, since most DI subunits were also observed unmodified. No monomeric DI was observed.

The remaining major families are CTL (14%), with the two previously mentioned VLFXA light chains as well as homologs to the CTL 3A, B9, A14, A15 and 4B, and PLA_2_ (13%). The venom contained only two PLA_2_ (13%), eluting around 80 min in the HPLC profile. They were identified as acidic phospholipase A2 1 (6.4%; *m/z* 13662.79, non-red.) and A2 2 (6.4%, *m/z* 13644.79, non-red.) from *M. lebetinus* and their sequences have been confirmed by TD between the reduced and non-reduced samples, including validation of their seven disulfide bridges, each. Additionally, LAAO (4%), CRISP (0.9%), NGF (0.8%) and PDE (0.2%) were detected as less dominant toxin families.

The venom profile of the analyzed *M. l. obtusa* is dominated by one peptide containing peak (F5), with 9% of the whole venom. It has two major molecular masses of *m/z* 444.22 (pEKW) and its 2M+H^+1^ ion of *m/z* 887.44. Further abundant peptides are pEKWPSPKVPP (*m/z* 1146.63) and pEKWPVPGPEIPP (*m/z* 1327.71).

### 3.4. Daboia palaestinae

The last Viperinae genus *Daboia* is represented by *D. palaestinae*. Our analysis revealed, that its venom is largely composed of svMP (22%) with only P-III svMP (16%) and DC proteins (6%), as well as an abundant amount of CTL (21%) (**Figure 5, Supplementary Table S9, S16, S23**). The earlier eluting CTL at R_t_ = 82 to 88 min (F28-33) have been annotated by several tryptic peptides as homologs to *M. lebetinus*, while the later (R_t_ >90 min) are related to Snaclec 3 and Snaclec 4 (*D. palaestinae*). The third abundant toxin family, svSP (18%), is described by different fibrinogenases and plasminogen activators. The HPLC venom profile lacks any dominant peak between R_t_ = 60 and 75 min and shows one abundant peak at R_t_<80 min (F26/27), which normally contain the PLA_2_ and CRISP variants in viper venoms. Therefore, no CRISP were observed and all PLA_2_ (8%) were described within F26/27 and being acidic. The two main proteoforms (*m/z* 13672.78 and *m/z* 13687.77) were TD identified as VpaPLA2 and VP7 from *D. palaestinae* and MVL-PLA2 from *M. l. transmediterranea*, since all three PLA_2_ sequences are highly identical with changes at only three amino acid positions.

Secondary toxin families in the venom of *D. palaestinae* are VEGF (11%), mainly homolog to VR-1 from *D. siamensis*, LAAO (2%) and KUN (2%). The ion mass of *m/z* 7722.582 was identical to then KUN serine protease inhibitor PIVL from *M. l. transmediterranea*, and to the its ^64^IQPR*^C^ C*-terminal shortened variant (*m/z* 7228.23). The only DI (0.5%) is the small KTS sequence containing viperistatin with *m/z* 4469.84 and four TD confirmed disulfide bridges. No minor or rare toxin families were observed within the Turkish *D. palaestinae* venom.

The peptidic part (14%) includes as main representatives, two svMP-i (pEKW, pENW) already detected within the other viper venoms of this study. But while no pERW mass has been observed, several related sequences could be annotated, such as pERWPGPKVPP (*m/z* 1144.63) and pERWPGPELPP (*m/z* 1159.59).

## 4. DISCUSSION

To gain better insights into the venom variations and the potential impact of medical significance of Palearctic vipers, we aligned the data of the seven vipers in a genus-wide comparison (**Figure 6**). For this purpose, we updated the previous venomics database of the full Old World viper subfamily (Viperinae) from Damm *et al.* (2021) and added additional snake venomics studies of Palearctic vipers until the end of 2023, searched by identical parameters ^16^.

**Figure 6.**
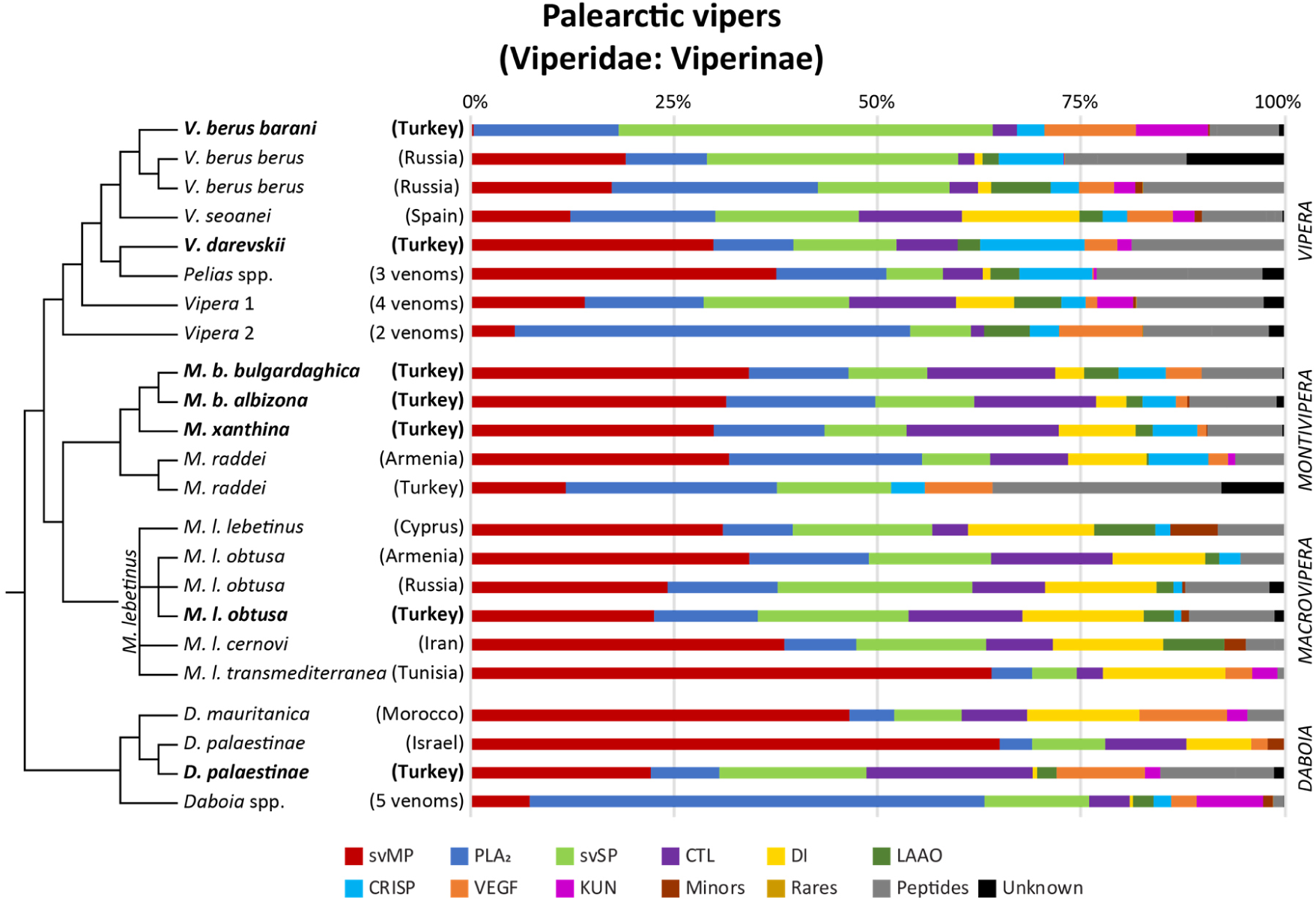
Snake venomics of Palearctic viper venom proteomes. Overview of all four genera (*Vipera*, *Montivipera*, *Macrovipera* and *Daboia*) summed to the total composition of Palearctic viper venoms (pie chart) and updated according to Damm *et al.* (2021). The 33 comparative proteomics data of 15 different Viperinae species including subspecies are lined up phylogeny-based. Origins of investigated specimen are reported in brackets. Numbers represent investigations of >1 venom proteomes. Venoms from this study are in bold. Schematic cladograms of the phylogenetic relationships based on Freitas *et al*. (2020).

### 4.1. *Vipera* - Eurasian vipers

With more than 20 species the Eurasian vipers (genus *Vipera*) are the most diverse group of all Old World vipers and can be split into three major clades ^18^. The *Pelias* group includes the common adder *V. berus* and meadow vipers of the *V. ursinii-renardi* complex. The other two groups in *Vipera* 1, comprising *Vipera aspis*, *Vipera latastei* and *Vipera monticola,* and *Vipera* 2, the nose-horned viper *V. ammodytes-meridionalis* complex, with its most recent changes ^88^. Several of those species are of medical relevance in Europe, i.a. *V. berus*, *V. ammodytes* and *V. aspis* ^15,89,90^. In Europe snakebite envenoming is an neglected health burden, even so over 5500 case have been reported in total ^89^. That said, in Europe no mandatory snakebite monitoring has been set up and therefore the true extent of snakebite envenoming remains to be clarified. Similar to the global situation, based on a combination of the non-mandatory snakebite reports and the lack of well-curated official databased statistics in most countries leading to high numbers of unreported cases, there are good reasons to assume that European snakebite incidences are certainly vast underestimated ^15,91^.

Above all, the adder *V. berus* with its extremely wide distribution is of particular interest for venom research, as it is still completely unknown to what extent a venom composition changes within a certain distribution range. Various factors such as genetic isolation and different habitats over several thousand kilometers across different climate zones with variable prey can have an unforeseen influence on the venom composition and make it impossible to predict variations ^29^. Therefore, it is surprising that relatively little is known about venom variations, both of nominal *V. berus berus* and the multitude of subspecies (*barani*, *bosniensis*, *nikolskii*, *marasso* and *sachalinensis*) ^10,92^. Only four venomic datasets (*V. b. berus* and *nikolskii*) have been reported beside our *V. b. barani* venom, with two Russian *V. b. berus* analysed by snake venomics ^16,93–96^.

Other studies over the past decades were based on single toxin isolation and characterization, or physiological effects ^92^. The two Russian *V. b. berus* snake venomics studies show the remarkable differences with the herein presented *V. b. barani* venom as svMP are nearly missing and is dominated by svSP, VEGF and KUN forming over 66% of the proteome (**Figure 6**). The only other *Vipera* descibed to harbor comparatively low svMP levels are *Vipera ammodytes montandoni* (1.8%) and the close related *V. b. nikolskii* (0.7%), sometimes also recognized as *Vipera nikolskii* ^16,62,96^. While high svSP contents are known for other Viperinae, like *Bitis* (15-26%), *Cerastes* (7-25%) or *Macrovipera* (5-24%) so far, only the venom of Russian *V. b. berus* with 30% svSP has been described with an increased svSP content ^16^. With 46% svSP the composition of the Turkish *V. b. barani* renders unique among so far quantified Old World viper venoms. Its most prominent protein, Nikobin, was firstly isolated from the *V. b. nikolskii* venom and is, like most svSP, a glycoprotein with unknown glycosylation pattern and putative hemotoxic activity ^97,98^. Sequences of the proteins show three *N*-glycosylation recognition sites, which high potential variability would explain the complex peak pattern observed for the *V. b. barani* venom profile. It is questionable to what extent the clinical manifestations would be similar, as there is only one suspected case report of this subspecies to date ^99^. In addition to local swelling, and hyperemia, there were clear neurological symptoms with pronounced diplopia and ptosis. No further symptoms were described after two ampules of antivenom (European viper venom antiserum, ‘Zagreb’). The bites of *V. berus* have a broad spectrum of potential effect, and is often per se defined as cyto- and hemotoxic with pro- or anticoagulant inducing effects and blood factor X activators ^92,100^. However, one problem is that the neurotoxic effects of *V. berus* envenoming are poorly documented in comparison to the amount of bite cases, but known for the other two medical relevant species, *V. aspis* and *V. ammodytes* ^24,101–105^. PLA_2_, such as presynaptic ammodytoxin isoforms and postsynaptic isoforms of aspin and vipoxin, are most likely responsible for these effects ^94,106,107^. This toxin family could be detected in all *V. berus* venom proteomes in varying abundances and the venoms of *V. b. nikolsksii* and the Slovakian *V. b. berus* were described being largely PLA_2_-rich, as many Russian vipers of the *Pelias* group ^16,96^. The impact of the extremely high svSP content in *V. b. barani* might be accompanied by strong effects on coagulation pathways and platelet aggregation like in other vipers ^98,108^. This shows that the venoms of the Eurasian adders are far more complex than previously investigated and thus represents an important subject for future venom research with a high relevance for the therapeutic treatment and specimen/population selection for antivenom development.

Within Europe several antivenoms are available with vipers as immunizing species. This include usually the four medically most important snakes *V. ammodytes*, *V. aspis*, *V. berus* and *V. latastei* ^89,90,100,109^. None of the antivenoms has been assessed by the WHO until now, but are registered by competent national authorities ^11^. A novel candidate with appropriate neutralizing potency is the polyvalent antivenom *Inoserp Europe* using seven species, including *M. xanthina* and two *Macrovipera* spp., with a broad cross-reactivity for European Viperinae ^110^. It needs to be noted, that many vipers of lower medical interest are often not tested and the antivenom efficiency against many of those taxa remains unknown ^111,112^.

Especially the taxonomically complex *Vipera* genus has several taxa with nearly no knowledge about bite consequences and their venom composition and pathophysiology ^9,18^. Their venom composition shows only a few rough trends of toxin family distribution as previously reviewed, whereby this complex picture has been further underpinned by more recent studies ^16,113,114^. Identified toxins within those neglected vipers often show homologies to highly active compounds of medically relevant taxa, such as *V. ammodytes* and *M. lebetinus*. One example is the here described *V. darevskii* venom, that is mainly dominated by svMP and confers to the classical Viperinae arrangement of major and secondary toxin families, like CRISP. Whether the described truncated *C*-terminal CRISP is an artificial cleavage product of the main toxins or an independently functional toxin cannot be determined from its sequence alone. Nevertheless, it is striking that it represents a self-contained and structurally stabile subdomain with five disulfide bridges, referred to as the Cysteine-Rich Domain (CRD) or Ion Channel Regulatory (ICR) domain ^115^. This domain contains the ShKT superfamily-like sequence known from highly potent small venom peptides produced by anemones with a strong effect on potassium channels ^116^. Similarly, in snake venoms other *C*-terminal subdomains are known to have evolved into independent toxins, such as DI and DC proteins from svMP ^117–119^.

Additionally, such neglected taxa have similar large proportion of peptides, consisting of BPP and natriuretic-related peptides, which even at low concentrations can have serious effects on the corresponding physiological systems. With high homology or even identical sequences to the BPP of pit vipers, as the most famous *Bothrops jararaca*, suggests that these peptides may also be responsible for corresponding responses in Palearctic vipers as herein described for all four genera, and discussed later in detail ^120^.

### 4.2. *Montivipera* - Mountain vipers

The mountain vipers (genus *Montivipera*) are divided into two clades, the Ottoman viper *M. xanthina* including *M. bulgardaghica* and the *M. raddei* complex. In comparison to the other three Palearctic viper genera, little is known about their venoms and the clinical consequences of a bite, since only a few studies report on *Montivipera* envenoming ^58,121,122^. Reported bites are from Türkiye, Armenia, Lebanon and Iran and describe symptoms reaching from local effects such as extensive blistering, local edema and necrosis up to coagulopathy and leucocytosis, and in two cases with lethal consequences ^121,123^.

Our mass spectrometric analysis revealed that the venoms of the three examined *Montivipera* spp. are relatively similar. A genus-wide comparison showed, that also the venom profile of the Armenian *M. raddei* has also a similar composition (**Figure 6**). The *M. raddei* venoms from Armenia and Türkiye are surprisingly divergent, and for the Turkish population only five toxin families have been reported. These include nearly 30% peptide content and 8% of unknown identity ^58,124^. Our discovery of PLA_2_, VEGF and CTL homologs to toxins of *D. russelii, D*. siamensis, *M. lebetinus* and *V. ammodytes* in all three *Montivipera* venoms emphasises their potential hazardous nature. The intravenious murine LD_50_ for Iranian *Montvipera latifii* and *M. xanthina* was determined to be <0.5 mg/kg, in the same range as Caspian cobra *Naja oxiana*, saw-scaled viper *Echis carinatus* and *M. lebetinus* (determined in µg venom per 16-18 mg mouse), analogous to the results of a comparison of 18 different Palearctic viper taxa ^110,125^. The similarities found for such snakes of medical relevance indicates that the genus *Montivipera* is of comparable danger. Consequently, bites must be treated with equal caution particularly at the hemo- and neurotoxic level. This is exemplified by several *Montivipera* spp. venoms with potent anticoagulant effects on human plasma ^126^. The WHO lists only a few antivenoms with *Montivipera* taxa as immunizing venom species, namely *M. xanthina* and *M. raddei*, including the previously mentioned Inoserp Europe ^11,15,110^. Therefore, it remains questionable whether such antivenoms are effective against the lesser known *Montivipera* species, especially since some venom are similar at the intra-genus level (here four of five proteomes), but can be strongly variable at the species level, like in *M. raddei* (**Figure 6**).

### 4.3. *Macrovipera* - Blunt-nosed vipers

The blunt-nosed vipers *Macrovipera* are widely distributed in the Middle East ^127,128^. Its most widespread representative, *M. lebetinus*, including several subspecies, can be found in over 20 countries and is by the WHO listed as highly medical relevant in more than half it ^11,21,22^. A detailed genus-wide comparison of all blunt-nose vipers venoms has been published recently in tandem with a detailed biochemical and pharmacological overview of *M. lebetinus* ssp. toxins ^129,130^. Thus, these aspects will only be briefly discussed here.

The overall composition of our Turkish *M. l. obtusa* venom mirrors that of the Armenian and Russian *M. l. obtusa*, and also the other subspecies (*M. l. lebetinus* and *cernovi*) share a similar compositions, with the *M. l. cernovi* venom showing the largest divergence (**Figure 6**). The taxonomically debated African subspecies *M. l. transmediterranea* is a clear outlier, with a noteworthy increased proportion of svMP. With its VEGF and KUN, the venom is more similar to *D. mauritanica*, which also occurs in the areas of North Africa. Furthermore, P-III svMP including DC proteins are beside svSP, the most prominent toxins across all *M. lebetinus* venoms. The CTL, partially forming the trimeric VLFXA complex with a P-III svMP, have higher variation (3-15%), similar to LAAO (0-8%). It should be emphasized that *Macrovipera* has the largest DI amount of the four genera with a consistently high content of 11-16%, independently to the DI subfamily composition. Although the expected monomeric, KTS sequence containing short DI obtustatin was originally characterized as high abundant toxin of *M. l. obtusa* (unreported local origin) with 7% of the whole venom proteome, no short nor monomeric DI has been described until now for any Turkish and Iranian *Macrovipera* venom ^129,131^, while several R/KTS DI are even known from other Viperidae venoms, including recently *Vipera* ^114,132^. Similarly, the venoms of another Turkish *M. l. obtusa* location and an Iranian *M. l. cernovi* lack small DI, while the Russian and Armenian *M. l. obtusa* contain them ^129^. This indicates that the subfamily of monomeric R/KTS DI is diversely distributed even within the genus *Macrovipera*. A detailed understanding of DI heterogeneity is of clinical importance and accordingly, this aspect demands further investigation in the future. A sequence clustering showed, that dimeric and short DI are the closest related snake venom DI subfamilies and might be a hint for this shift in their composition ^133^. A previous study, focusing on the Milos viper (*M. schweizeri,* recognized as a subspecies of *M. lebetinus* by several authors) and three *M. lebetinus* ssp. showed similar HPLC, SDS and bioactivity profiles ^129^. On the clinical side, it is therefore to be expected that the symptoms across the investigated *M. lebetinus* ssp. localities might be similar with effects on hypotension, hemorrhage and strong cytotoxicity leading to necrosis ^134,135^. On the other side, the geographic distribution of *Macrovipera* is large and includes an array of environments, so it is difficult or even impossible to predict venom variation, equal to the earlier mentioned *V. berus*. Such assumptions need to be investigated in the future through case reports or venom samples from different areas, as it has been done in recent years for the Indian Russell’s viper (*D. russelii*), for example, where initial generalizing assumptions led to serious complications in antivenom production and treatment^33,136^.

### 4.4. Daboia

The *Daboia* spp. ranks among the most medically significant snake lineages. They consists of a venom-wise understudied western Afro-Arabian group (*D. mauritania*, *D. palaestinae*), and the eastern Asian group, with *D. russelii* belonging to Indians ‘Big Four’. About 18 venom proteomes have been published for *D. russelii*, in addition to the eleven of the closely related *D. siamensis*, formerly *D. russelii siamensis* (**Supplementary Table S2**). *Daboia* is a prime example for the effect of biogeographic venom variation, with notable effects on the limited antivenom usability across an entire distribution area. ^33^. This underlines how, not only on a genus-wide, but also on intraspecific venom variations manifest into a problem of high therapeutically and scientific interest.

The venom of *D. palaestinae* has been investigated three times in a venomics context, of which one has been quantified by peak intensities of a shotgun approach and two by snake venomics, but at different wavelength (230 nm versus 214 nm this study) ^137,138^. The other two were of Israeli origin, while this study based on the recently described Turkish population. Even if not all three studies can be directly compared, the two snake venomics approaches (Israel, Türkiye in this study) show already considerable differences (**Figure 6**). While the Israeli sample, similar to the *D. mauritanica*, is dominated by svMP (65%) and contains a relevant amount of DI (8%), the Turkish venom shows a rather unusual composition, as previously described in detail. In particular, the lack of DI and the high level of VEGF distinguish it from the Israeli proteome from 2011 ^137^. The Israeli shotgun composition from 2022, on the other hand, even lists svSP as the main toxin group, followed by CTL and PLA_2_, while the svMP only make up a marginal proportion of the identified peptides (3%)^138^. With these different analytical methods in mind, it shows clearly that all three *D. palaestinae* venoms have a significantly different composition. While Laxme *et al*. (2022) reported in a direct comparison that the Israeli *D. palaestinae* is svSP and the Indian *D. russelii* svMP dominated, Damm *et al*. (2021) showed in a proteomic meta-analysis that *Daboia* venoms are more split into an Afro-Arabian and an Asian *Daboia* venom clade ^16,138^. They are dominant in SVMPs with DI in the western clade, while PLA_2_ rich in the eastern clade, in contrast to the *D. palaestinae-russelii* comparison carried out by Senji Laxme *et al*. (2022). However, the herein newly reported venom composition of the Turkish population does not exactly fit to either assignment. To what extent the venoms of *Daboia*, and *D. palaestinae* in particular, are really that multivariant or artifacts of different analysis methods needs to be clarified in future.

The bites of *D. paleastinae* are well studied for humans, but also in dogs, horses and further pets or farm animals ^139–142^. Due to the presence of the similar toxins in the here presented venom, it can be assumed that the clinical symptoms of envenoming by Turkish specimen are similar to those of other localities. No bites from the distant Turkish region are yet reported. Nevertheless, the different abundances of the toxin families could result in altered severity of the symptoms. A previous bioactivity-guided study on the hemotoxic properties revealed that *D. palaestinae* venom from different localities (twice Israel, once unknown) had evident variation in its activity across most of the tested assays ^143^. Especially the strongly reduced svMP and DI in the Turkish venom, as well as the increased proportion of svSP and VEGF might have severe influence on the degree of platelet aggregation and blood clotting.

### 4.5. Small venom peptides of Palearctic vipers

The proteomic landscapes of snake venoms are intensively investigated and reviewed ^16,81,144^. However, the knowledge about their lower molecular weight, peptidic compounds more restricted. While several of the larger peptide families, with sizes up to 9 kDa, are often reported as toxin families on their own (such as three-finger toxins (3FTx), KUN, DI or crotamine), components below 4 kDa are largely neglected ^145,146^. While a variety of bradykinin potentiating peptides (BBP), which were with their strong hypotension activity a template for Captopril, are known from Crotalinae venoms, only few studies looked into the peptidome of Viperinae ^16,120^.

Our rigorous MS profiling allowed us for the first time, to identify an array of low molecular weight peptidic components from the seven herein analyzed taxa. As mentioned in the previous part, i.e. KUN and different DI are well known for viperine venom and were usually identified in our analyzed samples. This indicates, that such peptides represent an important, yet seemingly often overlooked fraction of molecular diversity in viperine. While in *Vipera*, the peptide fraction fluctuated profoundly between taxa (ranging from 9-19%), the peptide landscape was more consistent in all three *Montivipera* spp. at 9-11%. *M. l. obtusa* and *D. palaestinae* showed 10-13%, respectively (**Figure 6**). Nevertheless, their compositions and the relative abundances of certain peptides differed strongly between the venoms and also within the same genera. Those identified peptides potentially originate from BPP and natriuretic peptide (NP) precursors, that can include repetitive svMP-i tripeptides and poly-His-poly-Gly (pHpG) sequences ^147^. A key element of most such peptides is the *N*-terminal pyroglutamate (pE), formed by glutaminyl cyclotransferases, which have been identified several times in viper venoms ^16^. The overall comparison showed strong similarities in the appearance of abundant peptides within *Montivipera*, the peptidome of which seems related to that of the *M. l. obtusa* (**Table 1**). Surprisingly, the peptidome of *V. b. barani* is more similar to *D. palaestinae*, than the taxonomically closer *V. darevskii*.

**Table 1.**
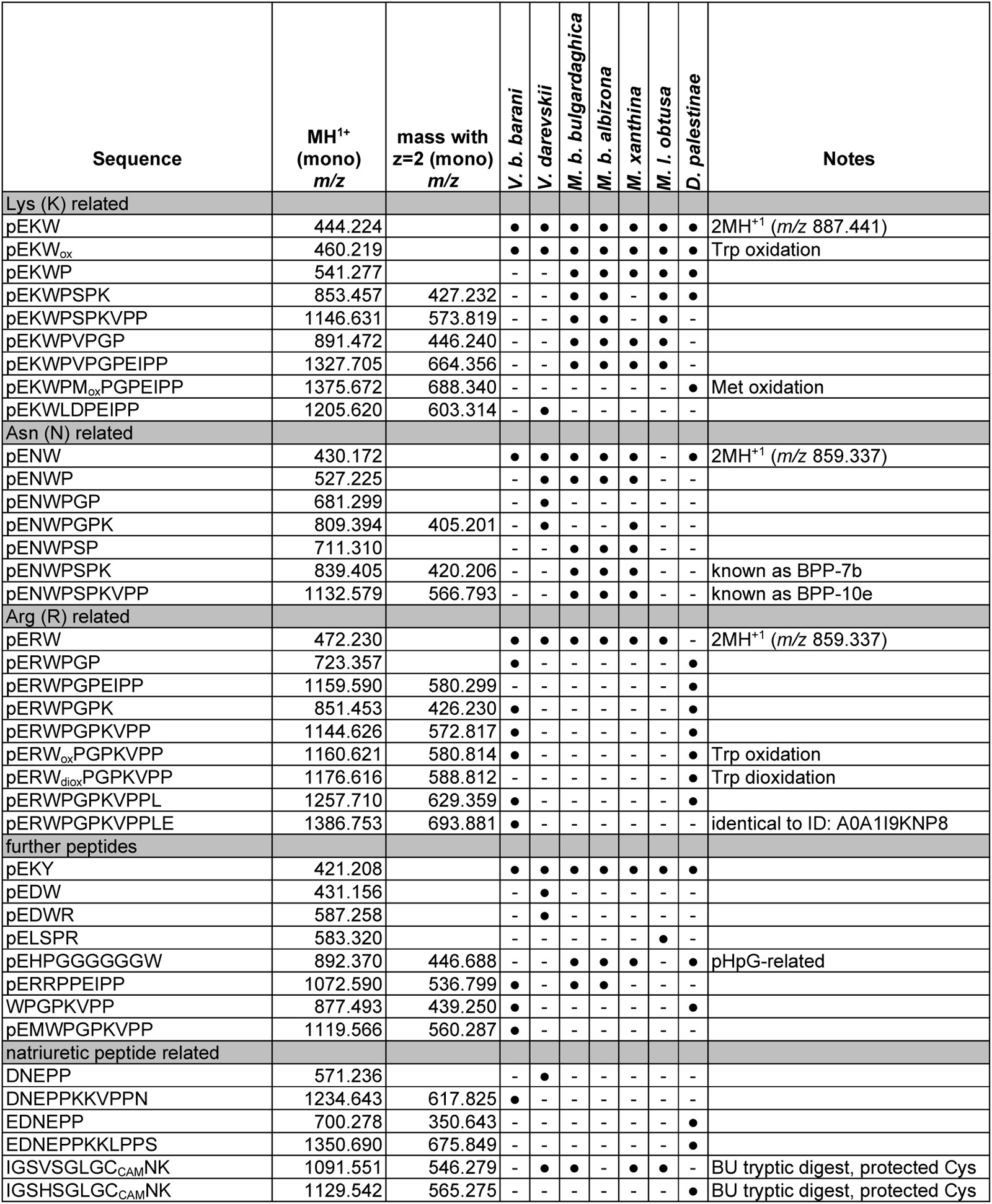
Peptidomics of svMP-i, BPP and NP of Palearctic vipers. Tandem MS/MS confirmed sequences of snake venom metalloproteinase inhibitors (svMP-i), bradykinin-potentiating peptides (BPP) and natriuretic peptides (NP) of seven viper venoms. Masses are given in monoisotopic (mono) *m/z* and if observed with double charges (z=2). Black dots mark the present of a peptide in the corresponding venom. Headline amino acid relation based on the modular pEXW, with pE for pyroglutamate and X for the mentioned amino acid. Amino acid I was set in similarities to known sequences, since a MS differentiation between isobaric L and I was not possible. Post-translational modification written out under ‘Notes’, as well as further information and carbamidomethyl (CAM).

Different BBP and *C*-terminal truncated sequences of variable length, from three to twelve amino acids, have been annotated in each of the viper venoms (**Table 1**). The shortest, tripeptidic sequences are henceforth referred to as svMP-i. These small peptides are predicted to protect the venom from auto-digestion by its own svMP ^148,149^. The three svMP-i (pEKW, pENW, pERW) are highly abundant, with pEKW often as main representative, and were detected in all seven venoms, except pENW, that could not be observed in the *M. xanthina* venom, and pERW in the *D. palaestinae* proteome.

Among the >25 oberserved peptides pEKWPVPGPEIPP was in all three *Montivipera* and the *M. l. obtusa* venom the main BPP-related sequence with Lys in second position and for the Asn-related pENWPSPKVPP (known as BPP-10e) is exclusive for *Montivipera* and pENWPGPK for *V. darevskii*. The Arg-related BPP were only abundant in the venoms of *V. b. barani* and *D. palaestinae* with various truncations of pERWPGPKVPPLE in both and pERWPGPEIPP in *D. palaestinae* only. The twelve-mer pERWPGPKVPPLE is identical to a building block of a *V. ammodytes* BPP-NP precursor (ID: A0A1I9KNP8_VIPAA) and a *V. aspis* BBP (ID: P31351). Additionally, Trp oxidations have been detected, like pEKW_ox_ in all seven venoms and a Met oxidation in pEKWPM_ox_PGPEIPP within *D. palaestinae*. Based on our observation, the BPP in Viperinae venoms following the modular structure of **pEXW(PZ)_1-2_P(EI)/(KV)PPLE**, with X mainly K/N/R, while other amino acids on position 2 are rare, Z = G/S/V and multiple *C*-terminal truncation. Some exclusive sequences, like the pEKWLDPEIPP (*V. darevskii*), pELSPR (*M. l. obtusa*) and pERRPPEIPP (*Vipera* and *Montivipera*), underlines that the whole group of BPP-NP precursor related peptides have a highly variable combination pattern, of which most physiological effects are still unknown. The high similarity to pit viper BPP sequences, suggests similar serious activities on the blood pressure.

The NP are the third group of peptides deriving from the same precursor. They strongly contribute to the lowering of blood pressure by the NP receptors via cGMP-mediated signaling. NP and can be found in various animals as well as the venom of some elapids and vipers ^150^. Snake venom NP are structurally homolog to mammalian NPs, including the conserved 17-residue ring structure, closed by a disulfide bridge, with an *N*- and *C*-tail region of variable length ^151^. Their molecular size ranges from 2-4 kDa and they are known from highly medical relevant snakes, like taipans (*Oxyuranus*), brown snakes (*Pseudonaja*), kraits (*Bungarus*) and blunt-nosed vipers (*Macrovipera*). In the case of *M. lebetinus* two different NP structures has been described as lebetins: the long lebetin 2 (3943.4 Da, with one disulfide bridge) and the short lebetin 1 (1305.5 Da), which is identical to the lebetin 2 *N*-terminus ^152^. This terminal sequence is known to be important for platelet aggregation inhibition and to prevent collagen-induced thrombocytopenia^153^. We observed two peptides with sequences similar to the short lebetin 1β (DNKPPKKGPPNG), those are DNEPPKKVPPN in *Vipera* with K2E and G8V, as well as EDNEPPKKLPPS in *Daboia* with an additional *N*-terminal Glu and three substitutions (K2E, G8L and N11S) (**Table 1**). The longer lebetins were full length detected in the venom of *M. l. obtusa* as expected for a *M. lebetinus* subspecies, but surprisingly also in *M. b. bulgardaghica* with a homolog to lebetin 2α. Further tryptic peptides of NP related sequences, has been observed in *V. darevskii* (gel band 12a), *M. b. bulgardaghica* (16a), *M. xanthina* (10a), *M. l. obtusa* (8a). For example, all genera showed the *C*-terminal IGSVSGLGCNK sequence, with a single amino acid change of H4V, except *Macrovipera*, that had the lebetin 2 identical *C*-terminal sequence of IGSHSGLGCNK. Therefore, we confirmed the appearance of NP in the venom of all four genera at the proteomics level, which seems to be a constant part of Viperinae venoms in general.

## 5. SUMMARY

Palearctic vipers are a highly diverse group of venomous snakes with high impact on health and socioeconomic factors that can be found across three continent. By extensive venomics studies on seven different taxa from Türkiye within this group, the venom proteome and peptidome was characterized and quantified in detail. Our complementary MS-based workflows revealed high divergence in their abundance of toxin families, following the major, secondary and minor toxin family trend known for Old World vipers. A closer look into the type of toxins and corresponding abundancies shows notable differences between the investigated genera of *Vipera*, *Montivipera*, *Macrovipera* and *Daboia*.

Within the genus *Vipera*, *V. b. barani* had a unique venom mostly composed of svSP. This sets it clearly apart from *V. berus* venoms of other localities, but also viperine venoms in general. *V. b. barani* lacks svMP and the peptidome is closer to the highly medical relevant *D. palaestinae* than to the other viper venoms investigated in this study. Hence, the venom composition of *V. berus* cannot be easily generalized and in regard to its wide distribution and snakebite envenoming potential needs to be more closely investigated in future studies. The venom of *V. darevskii*, as an example of an understudied taxa, which was unknown until now. We could show, that its composition based on different myotoxic and anticoagulant active homologs, as well as a highly abundant pEKW peptide part of >10% of the total venom composition. Furthermore, within its venom a truncated but presumably self-contained *C*-terminal CRISP subdomain could be annotated. It includes a ShKT-like, or CRD domain, indicating potential neurological envenoming effects by *V. darevskii*. Beside, the parallels between our *V. b. barani* and *D. palaestinae* venom, we could also show important similarities within the genera *Montivipera* and *Macrovipera* on both, proteomics and peptidomics, level. Here, we describe the first genus-wide *Montivipera* venom comparison. The venom compositions across four taxa of the subclades *raddei* and *xanthina* have a consistent appearance, with the Turkish *M. raddei* as an outliner until now. The direct comparison of the three *Montivipera* venom profiles consistently showed a wide range of toxin homologs to highly medical relevant viper species.

The herein investigated venom of *D. palaestinae* is in support of of a high venom varation within the genus *Daboia*. As it is known for eastern *Daboia* species to cause locality-based different clinical images after a bite, we could show that also the western taxa have strong compositional differences. The *D. palaestinae* venoms of Türkiye and Israel display different toxin abundancies. Therefore, based on our findings it seems reasonable to expect that a high venom diversity like in Indian *D. russelii* might also be therapeutically relevant for *D. palaestinae*, if not even the whole genus *Daboia*.

Beside the well studied toxin families, all here investigated Palearctic viper venoms have a peptide content of at least 9%. They include a spectrum of svMP-i, BPP, pHpG and NP. We identified the modular consensus sequence **pEXW(PZ)_1-2_P(EI)/(KV)PPLE** for BPP related peptides in viper venoms. This underscores the intricate nature of snake venom peptidic compounds potential influencing blood pressure. Notably, they exhibit an increased impact on the venom composition, as evidenced by their prevalence not only in our seven vipers but also across various other viper species. Peptides found to be distributed in high proportions, equal to major toxin families, and, intriguingly, reaching even higher concentrations based on the small molecular weight. This points to the significance of BPP as well as NP in the overall venom composition, highlighting their potential role in the physiological effects following a snakebite envenomation, but might be often overlooked until now.

The study of the herein investigated seven Palearctic viper venoms shows, that their venoms include a variety of different potent toxin families. Since the vipers in Türkiye are responsible for numerous hospitalization of adults as well as children across the country each year, deciphering these venom variations is of great interest. Our data on the detailed venom compositions and the comparison to other proteomes, will contribute to provide novel biochemically and evolutionary insights in Old World viper venoms and emphasize the potential medical importance of neglected taxa. In particular, the first venom descriptions of several Turkish viper taxa, will facilitate the risk assessment of snakebite envenoming by these vipers and aid in predicting the venoms pathophysiology and clinical treatments.

## Supporting information

02a_SupportingInformation_Figures

02b_SupportingInformation_Tables

## AUTHOR INFORMATION

### Author Contributions

The manuscript was written through contributions of all authors. All authors have given approval to the final version of the manuscript. CRediT Taxonomy: Maik Damm (Conceptualization, Data Curation, Formal Analysis, Investigation, Project Administration, Visualization, Writing – Original Draft Preparation, Writing – Review & Editing); Mert Karış (Resources – Field work & Venom Milking, Writing – Review & Editing); Daniel Petras (Resources – Top-Down MS Measurements, Writing – Review & Editing); Ayse Nalbantsoy (Resources – Field work & Venom Milking); Bayram Göçmen (Resources – Field work & Venom Milking); Roderich D. Süssmuth (Funding Acquisition, Resources, Writing – Review & Editing).

### Notes

The authors declare no competing financial interest.

## ACKNOWLEDGMENT

We thank Tim Lüddecke, Ignazio Avella and Lennart Schulte for critical feedback and reviewing the early manuscript. We dedicate this paper to the memory of Prof. Dr. Bayram Göçmen, who lost his, fight against cancer. He was an outstanding teacher, a good friend and colleague, and loved by his family.

## ABBREVIATIONS

ABC: ammonium hydrogen carbonate
ACN: acetonitrile
BPP: bradykinin-potentiating peptides
CTL: C-type lectin-related proteins and snake venom C-type lectins
CRISP: cysteine-rich secretory proteins
DAD: diode array detector
DC: disintegrin-like/cysteine-rich proteins
DI: disintegrins
DTT: dithiothreitol
HFo: formic acid
KUN: Kunitz-type inhibitors
LAAO: L-amino acid oxidases
MES: 2-(N-morpholino)ethane sulfonic acid
NGF: nerve growth factors
NP: natriuretic peptides
PDE: phosphodiesterases
pE: pyroglutamate
pHpG: poly-His-poly-Gly
PLA_2_: snake venom phospholipases
A_2_: SDS, sodium dodecyl sulfate
svMP: snake venom metalloproteinases
svMP-i: snake venom metalloproteinase inhibitors
svSP: snake venom serine proteases
VEGF: vascular endothelial growth factors F
5N: 5′-nucleotidases.

---

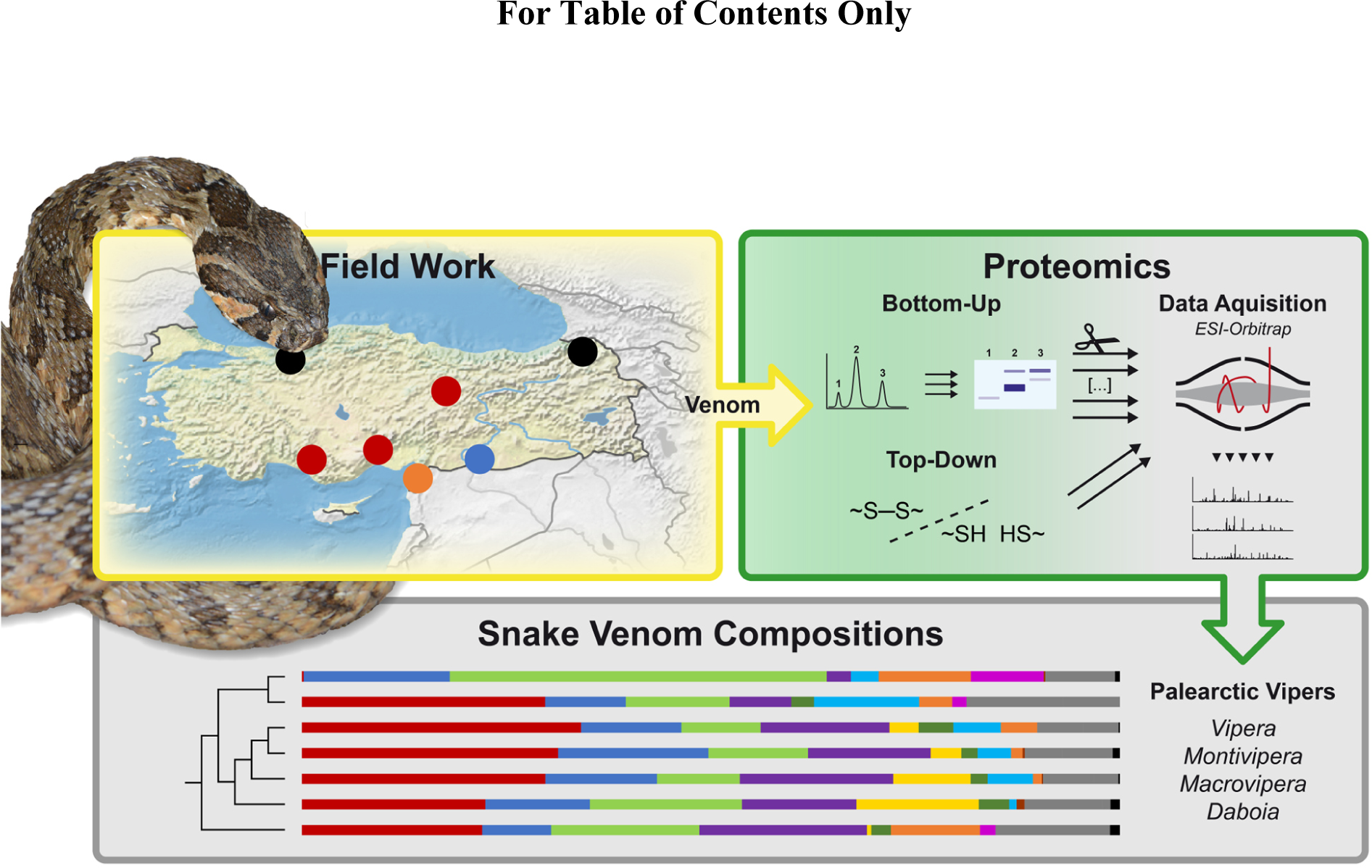

