## Supplementary material for "Venomics and Peptidomics of Palearctic vipers: Clade-wide analysis of seven taxa of the genera *Vipera*, *Montivipera*, *Macrovipera* and *Daboia* across Türkiye": 02a_SupportingInformation_Figures

Detailed supplementary tables are in the corresponding xlsx file.

**Supplementary Table S1: Venom pool information of the seven Palearctic viper venoms.**

**Supplementary Table S2: Database of Palearctic viper proteomes.**
All proteomes of Viperinae subfamily until 31st December 2020 are alphabetically listed by their current taxonomic status. The original taxon of publications is mentioned. Additionally, details about the used venom pool are listed, like origin, age and sex of the milked snakes, as well as the used methods for analysis and quantification for each proteome. Detected toxin families and, if quantified, their percentual values or general observation in a venom are given, with a detailed annotation of detected minor and rare families. Additionally, all publications used for main text Figure 1 are mentioned here from column AF ongoing, based on literature search from 2022 to 2023 as described, with earlier proteomes taken from the Old World viper venom proteome database (Damm et al. 2021).

**Supplementary Table S3: Quantification of the *V. b. barani* venom pool proteome.**
Fraction numbers are based on the RP-HPLC chromatogram. Band IDs are based on the SDS-PAGE profile. IMP masses and TD annotations are listed on the corresponding fraction.

**Supplementary Table S4: Quantification of the *V. darevskii* venom pool proteome.**
Fraction numbers are based on the RP-HPLC chromatogram. Band IDs are based on the SDS-PAGE profile. IMP masses and TD annotations are listed on the corresponding fraction.

**Supplementary Table S5: Quantification of the *M. b. bulgardaghica* venom pool proteome.**
Fraction numbers are based on the RP-HPLC chromatogram. Band IDs are based on the SDS-PAGE profile. IMP masses and TD annotations are listed on the corresponding fraction.

**Supplementary Table S6: Quantification of the *M. b. albizona* venom pool proteome.**
Fraction numbers are based on the RP-HPLC chromatogram. Band IDs are based on the SDS-PAGE profile. IMP masses and TD annotations are listed on the corresponding fraction.

**Supplementary Table S7: Quantification of the *M. xanthina* venom pool proteome.**
Fraction numbers are based on the RP-HPLC chromatogram. Band IDs are based on the SDS-PAGE profile. IMP masses and TD annotations are listed on the corresponding fraction.

**Supplementary Table S8: Quantification of the *M. l. obtusa* venom pool proteome.**
Fraction numbers are based on the RP-HPLC chromatogram. Band IDs are based on the SDS-PAGE profile. IMP masses and TD annotations are listed on the corresponding fraction.

**Supplementary Table S9: Quantification of the *D. palaestinae* venom pool proteome.**
Fraction numbers are based on the RP-HPLC chromatogram. Band IDs are based on the SDS-PAGE profile. IMP masses and TD annotations are listed on the corresponding fraction.

**Supplementary Table S10: Snake Venomics annotation of the *V. b. barani* venom by pFind.**
Toxin family annotation of tryptic digested single bands by pFind and BLAST.

**Supplementary Table S11: Snake Venomics annotation of the *V. darevskii* venom by pFind.**
Toxin family annotation of tryptic digested single bands by pFind and BLAST.

**Supplementary Table S12: Snake Venomics annotation of the *M. b. bulgardaghica* venom by pFind.**
Toxin family annotation of tryptic digested single bands by pFind and BLAST.

**Supplementary Table S13: Snake Venomics annotation of the *M. b. albizona* venom by pFind.**
Toxin family annotation of tryptic digested single bands by pFind and BLAST.

**Supplementary Table S14: Snake Venomics annotation of the *M- xanthina* venom by pFind.**
Toxin family annotation of tryptic digested single bands by pFind and BLAST.

**Supplementary Table S15: Snake Venomics annotation of the *M. l. obtusa* venom by pFind.**
Toxin family annotation of tryptic digested single bands by pFind and BLAST.

**Supplementary Table S16: Snake Venomics annotation of the *D. palaestinae* venom by pFind.**
Toxin family annotation of tryptic digested single bands by pFind and BLAST.

**Supplementary Table S17: Top-down annotation of the *V. b. barani* venom by TopPIC.**
Toxin annotation of non-reduced (grey) and reduced (no highlight) samples.

**Supplementary Table S18: Top-down annotation of the *V. darevskii* venom by TopPIC.**
Toxin annotation of non-reduced (grey) and reduced (no highlight) samples.

**Supplementary Table S19: Top-down annotation of the *M. b. bulgardaghica* venom by TopPIC.**
Toxin annotation of non-reduced (grey) and reduced (no highlight) samples.

**Supplementary Table S20: Top-down annotation of the *M. b. albizona* venom by TopPIC.**
Toxin annotation of non-reduced (grey) and reduced (no highlight) samples.

**Supplementary Table S21: Top-down annotation of the *M. xanthina* venom by TopPIC.**
Toxin annotation of non-reduced (grey) and reduced (no highlight) samples.

**Supplementary Table S22: Top-down annotation of the *M. l. obtusa* venom by TopPIC.**
Toxin annotation of non-reduced (grey) and reduced (no highlight) samples.

**Supplementary Table S23: Top-down annotation of the *D. palaestinae* venom by TopPIC.**
Toxin annotation of non-reduced (grey) and reduced (no highlight) samples.

**Supplementary Table S24: Dimeric disintegrins in the *M. l. obtusa* venom.**
Pairing and variation of disintegrins dimers identified by IMP and TD.

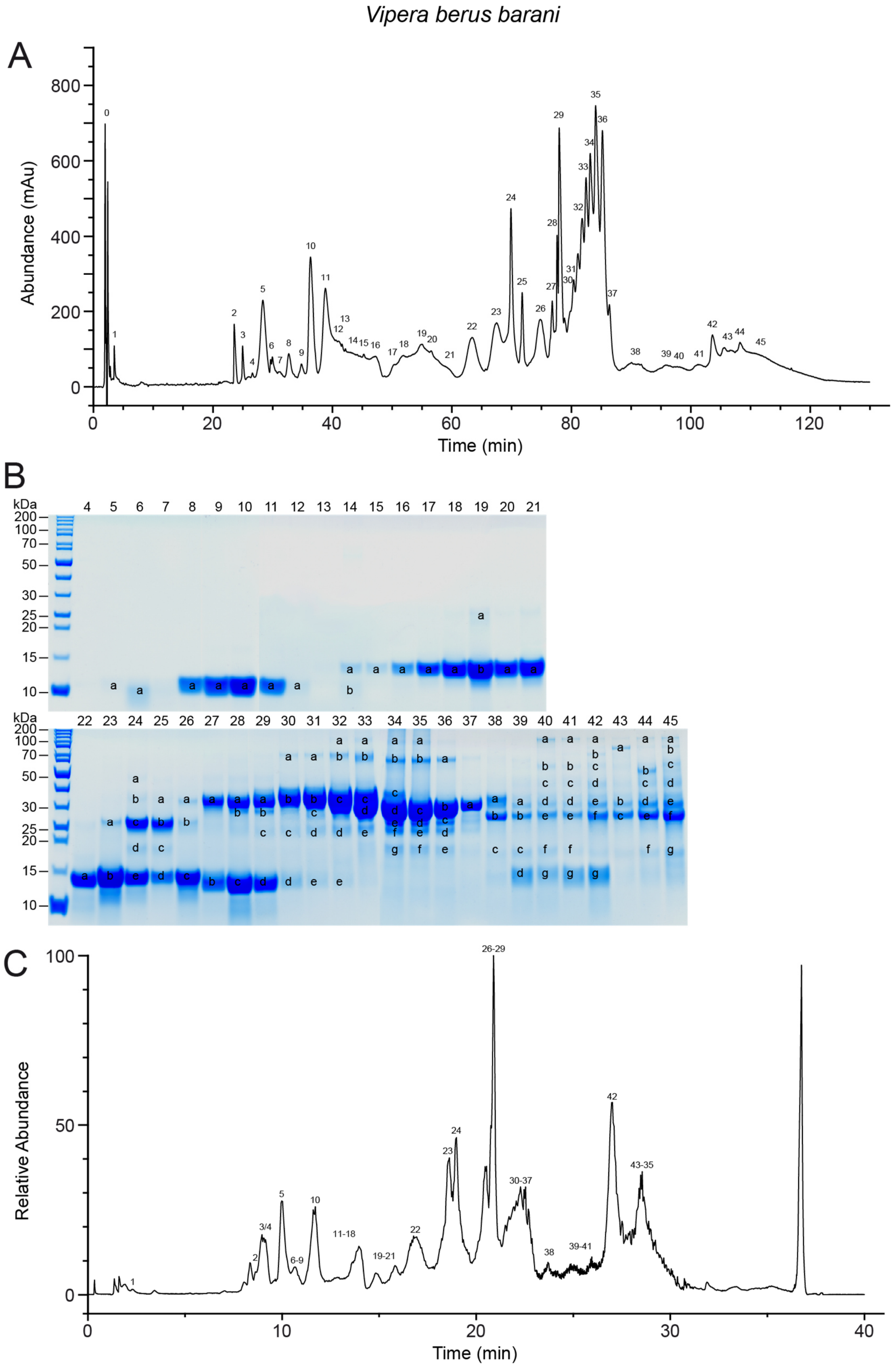


**Supplementary Figure S1: Venom profile of *Vipera berus barani*.**
For the bottom-up snake venomics approach the **(A)** RP-HPLC profile was observed at λ = 214 nm, with peak 0 corresponding to the injection peak, and venom fractions further analyzed by **(B)** Coomassie-stained SDS-PAGE under reducing conditions. PAGE line nomenclature is based on RP-HPLC fractions from (A). Labelled bands were cut, subjected to tryptic digestion, and analyzed with LC-MS. For top-down and intact mass profiling the venom profile was observed via the **(C)** total ion current chromatogram by ESI under non-reducing conditions.

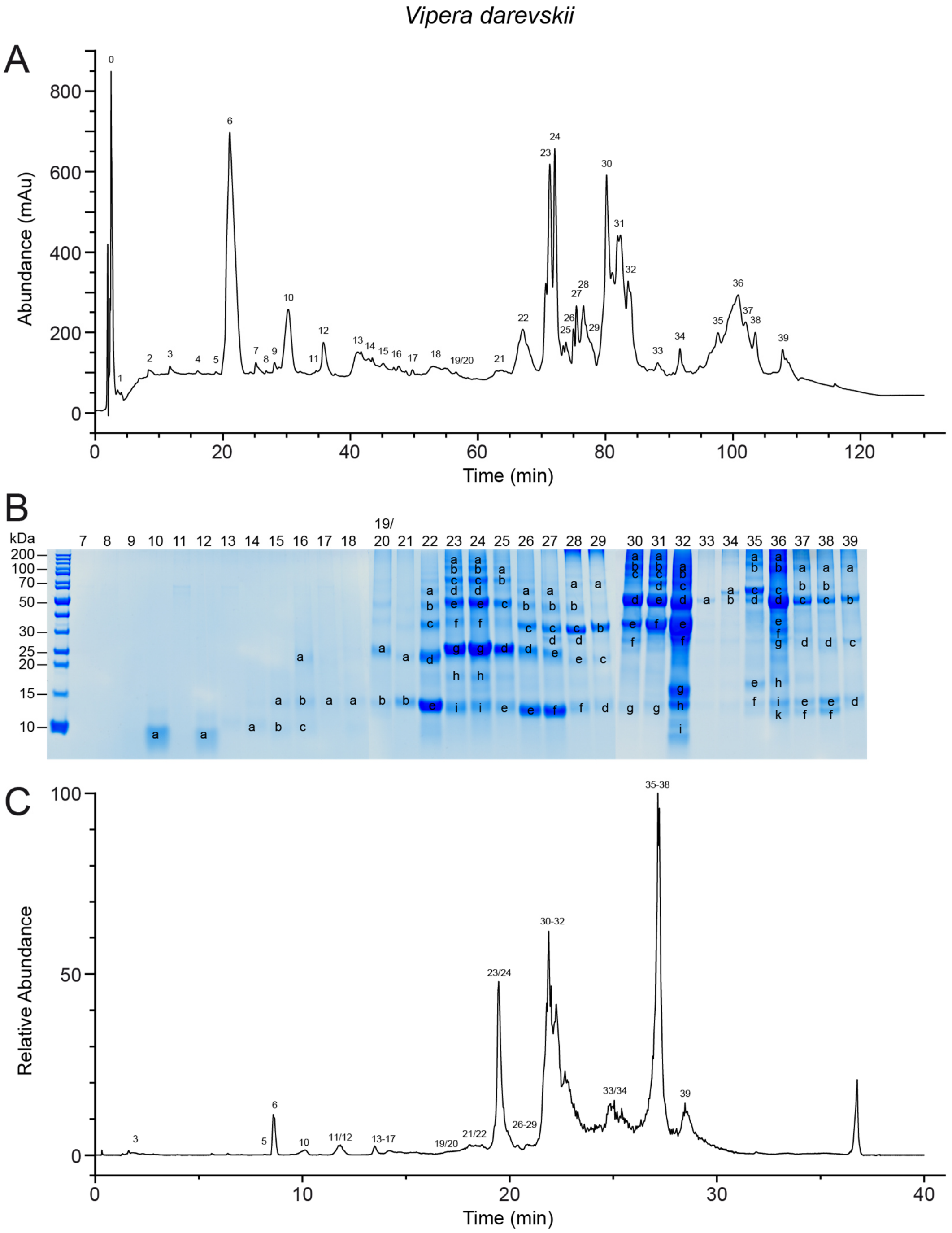


**Supplementary Figure S2: Venom profile of *Vipera darevskii*.**
For the bottom-up snake venomics approach the **(A)** RP-HPLC profile was observed at λ = 214 nm, with peak 0 corresponding to the injection peak, and venom fractions further analyzed by **(B)** Coomassie-stained SDS-PAGE under reducing conditions. PAGE line nomenclature is based on RP-HPLC fractions from (A). Labelled bands were cut, subjected to tryptic digestion, and analyzed with LC-MS. For top-down and intact mass profiling the venom profile was observed via the **(C)** total ion current chromatogram by ESI under non-reducing conditions.

*Montivipera bulgardaghica bulgardaghica*

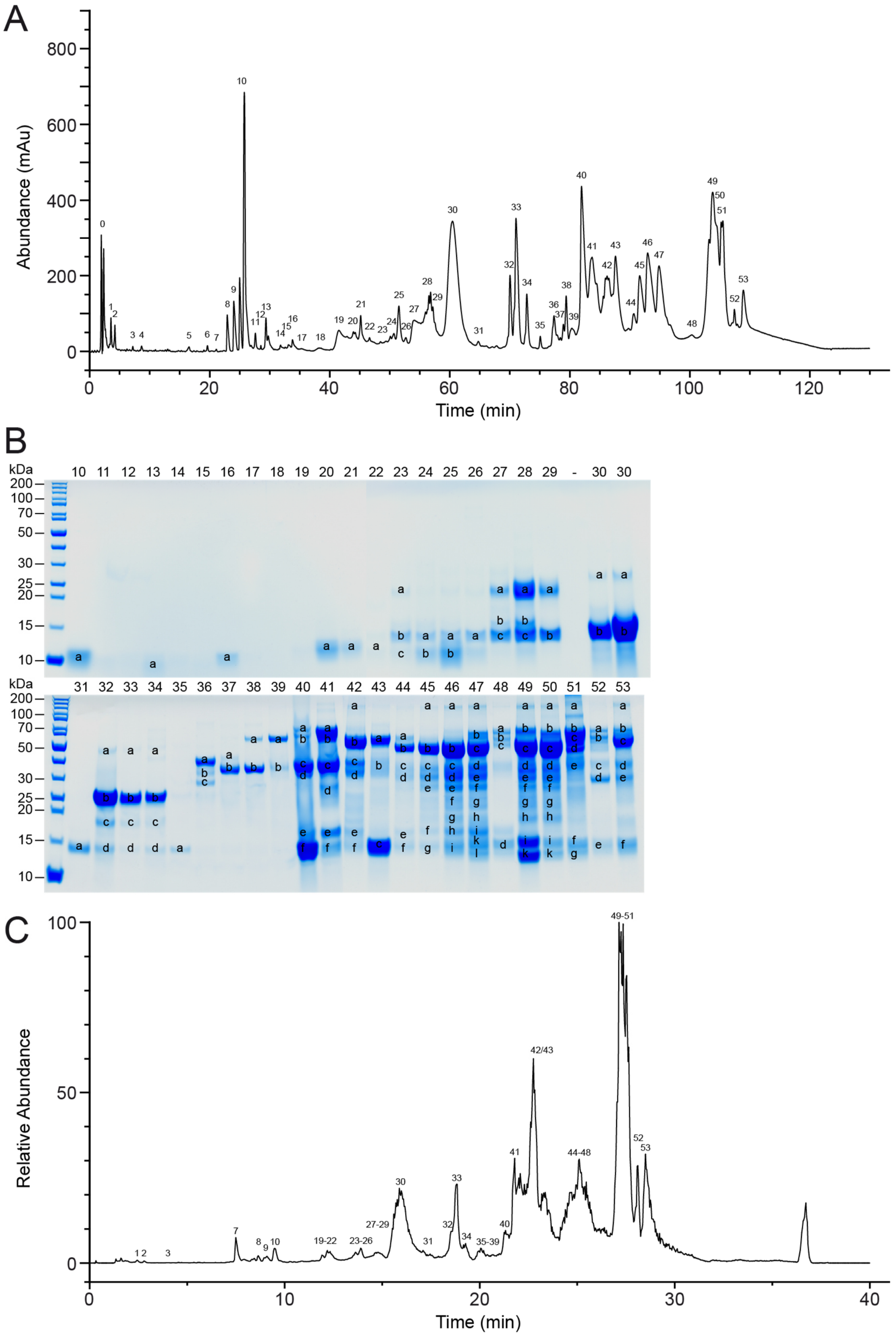


**Supplementary Figure S3: Venom profile of *Montivipera bulgardaghica bulgardaghica*.**
For the bottom-up snake venomics approach the **(A)** RP-HPLC profile was observed at λ = 214 nm, with peak 0 corresponding to the injection peak, and venom fractions further analyzed by **(B)** Coomassie-stained SDS-PAGE under reducing conditions. PAGE line nomenclature is based on RP-HPLC fractions from (A). Labelled bands were cut, subjected to tryptic digestion, and analyzed with LC-MS. For top-down and intact mass profiling the venom profile was observed via the **(C)** total ion current chromatogram by ESI under non-reducing conditions.

*Montivipera bulgardaghica albizona*

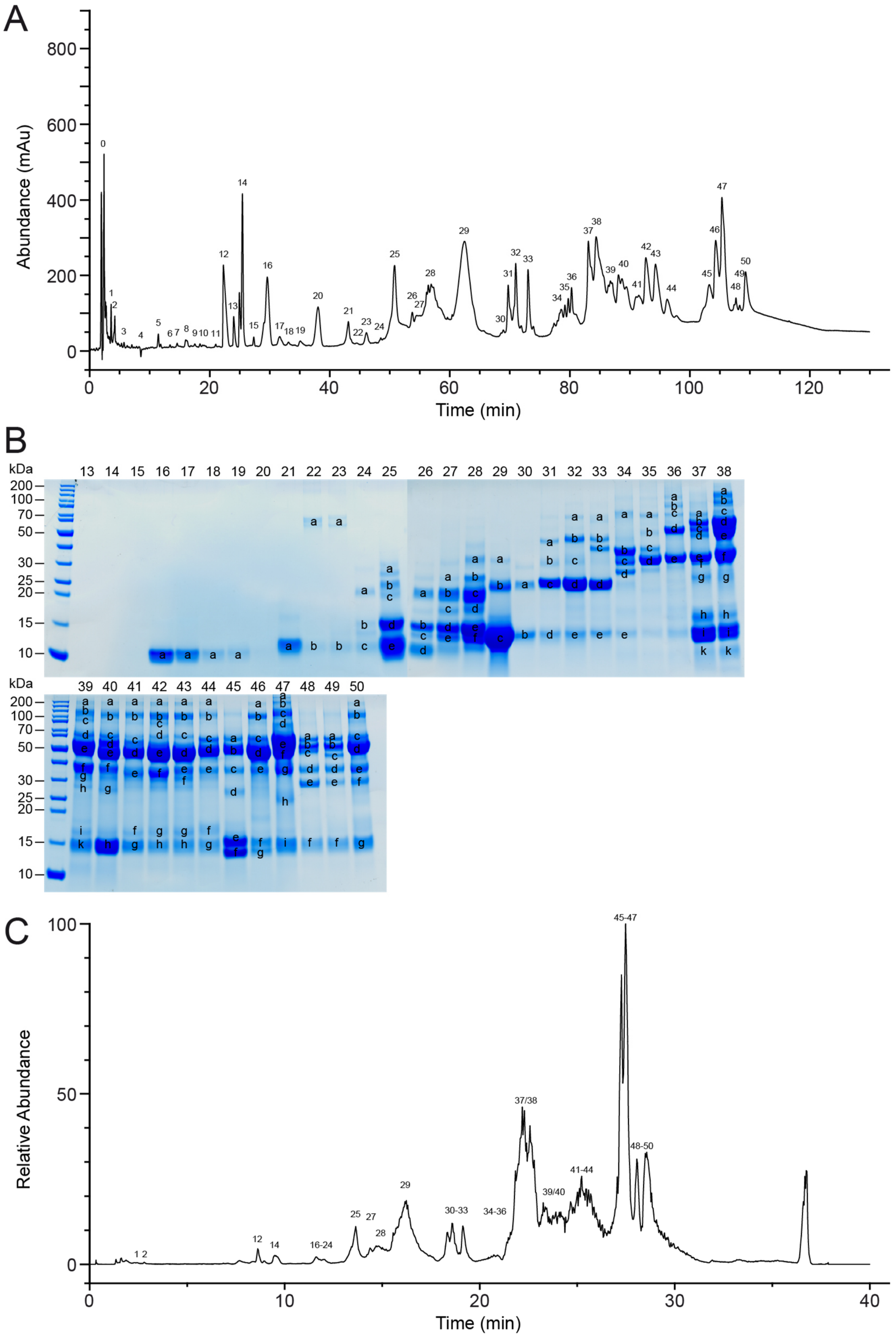


**Supplementary Figure S4: Venom profile of *Montivipera bulgardaghica albizona*.**
For the bottom-up snake venomics approach the **(A)** RP-HPLC profile was observed at λ = 214 nm, with peak 0 corresponding to the injection peak, and venom fractions further analyzed by **(B)** Coomassie-stained SDS-PAGE under reducing conditions. PAGE line nomenclature is based on RP-HPLC fractions from (A). Labelled bands were cut, subjected to tryptic digestion, and analyzed with LC-MS. For top-down and intact mass profiling the venom profile was observed via the **(C)** total ion current chromatogram by ESI under non-reducing conditions.

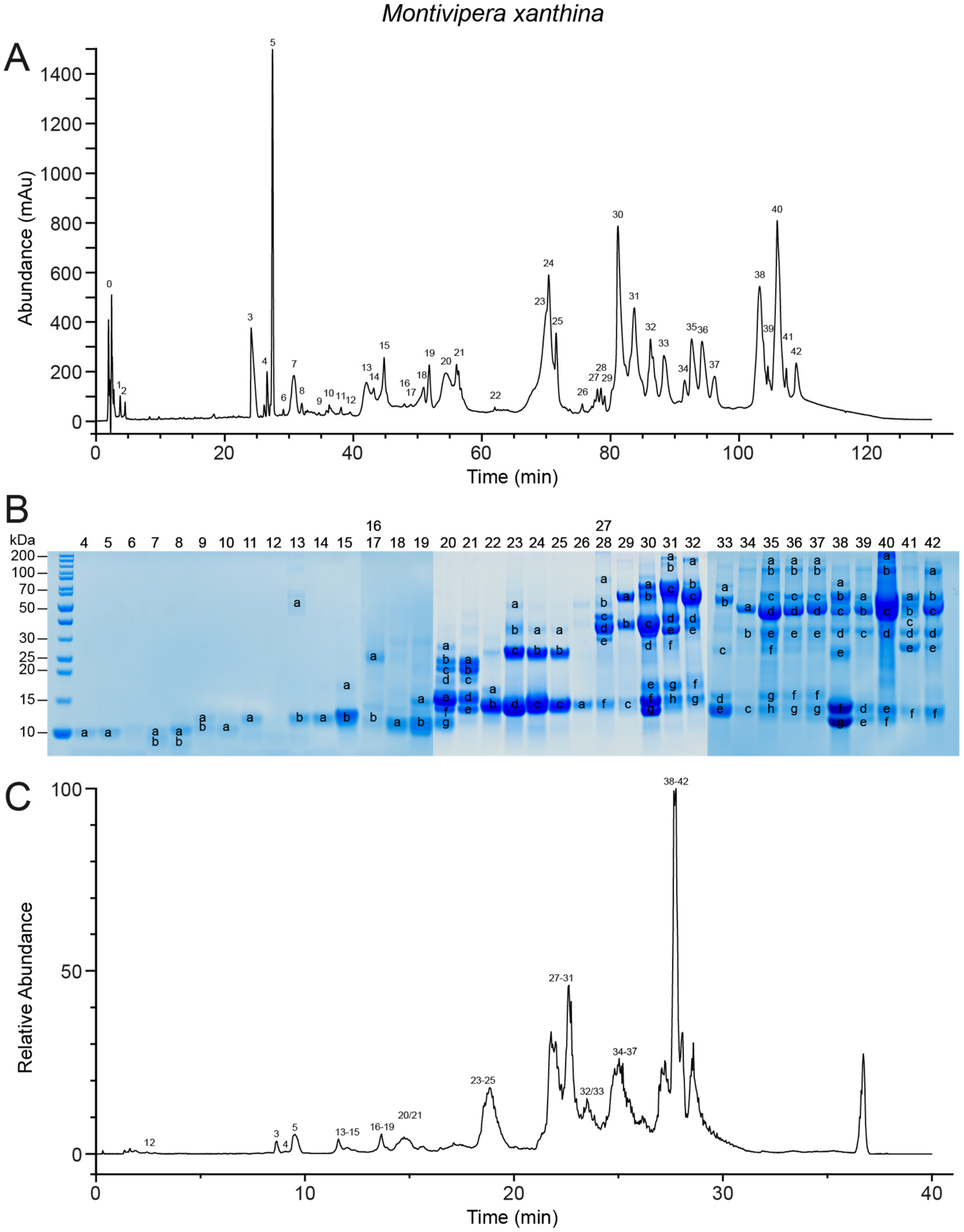


**Supplementary Figure S5: Venom profile of *Montivipera xanthina*.**
For the bottom-up snake venomics approach the **(A)** RP-HPLC profile was observed at λ = 214 nm, with peak 0 corresponding to the injection peak, and venom fractions further analyzed by **(B)** Coomassie-stained SDS-PAGE under reducing conditions. PAGE line nomenclature is based on RP-HPLC fractions from (A). Labelled bands were cut, subjected to tryptic digestion, and analyzed with LC-MS. For top-down and intact mass profiling the venom profile was observed via the **(C)** total ion current chromatogram by ESI under non-reducing conditions.

*Macrovipera lebetinus obtusa*

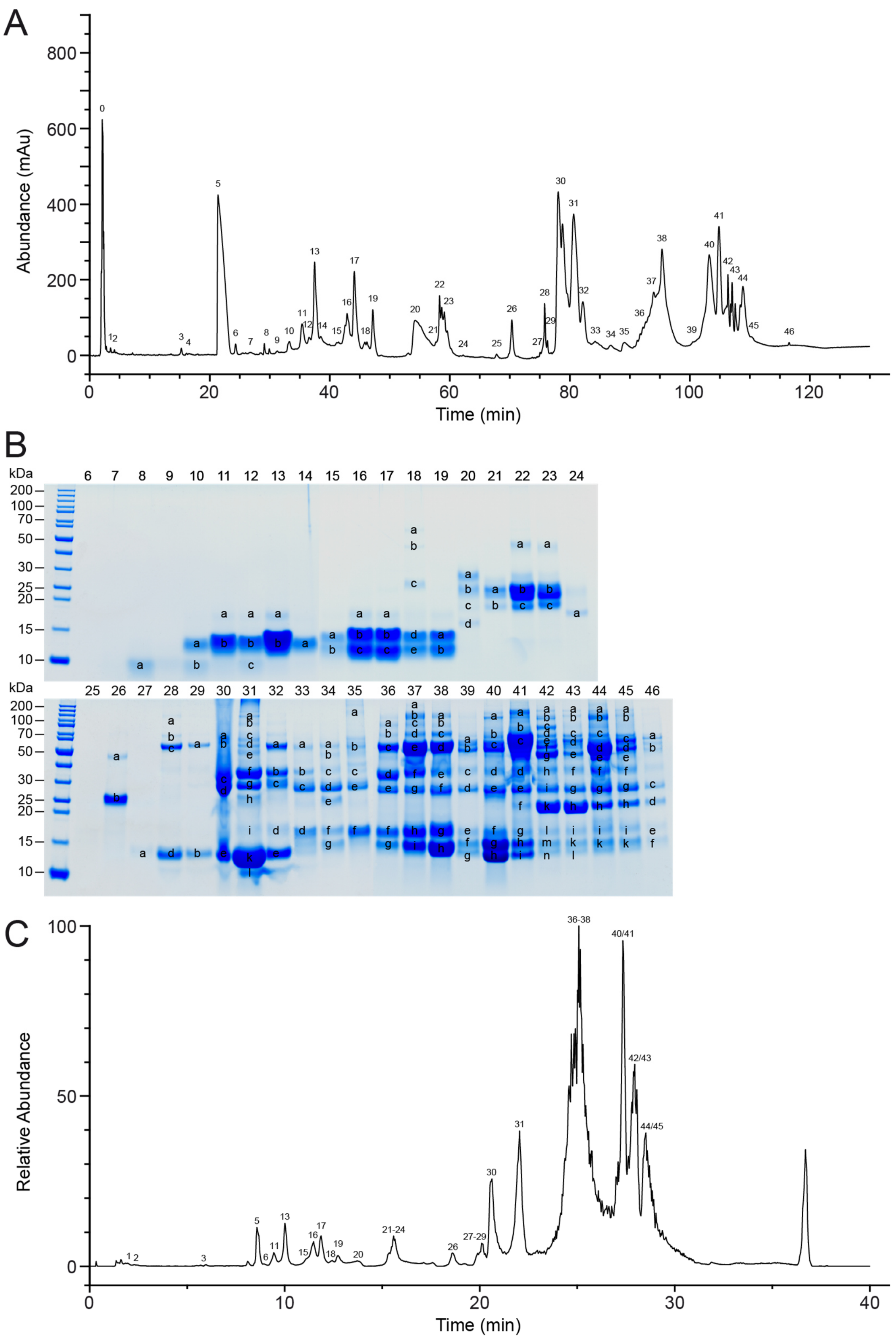


**Supplementary Figure S6: Venom profile of *Macrovipera lebetina obtusa*.**
For the bottom-up snake venomics approach the **(A)** RP-HPLC profile was observed at λ = 214 nm, with peak 0 corresponding to the injection peak, and venom fractions further analyzed by **(B)** Coomassie-stained SDS-PAGE under reducing conditions. PAGE line nomenclature is based on RP-HPLC fractions from (A). Labelled bands were cut, subjected to tryptic digestion, and analyzed with LC-MS. For top-down and intact mass profiling the venom profile was observed via the **(C)** total ion current chromatogram by ESI under non-reducing conditions.

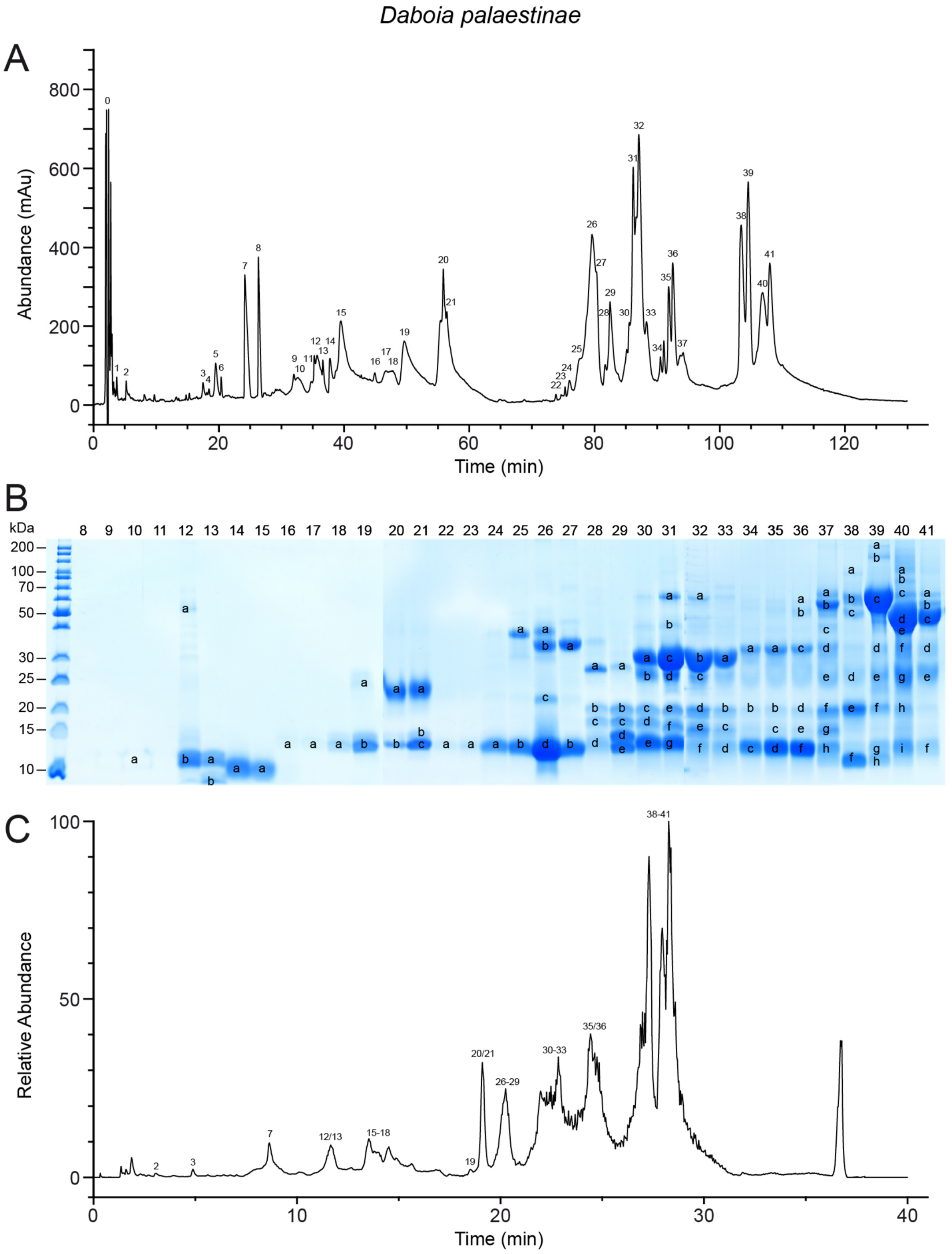


**Supplementary Figure S7: Venom profile of *Daboia palaestinae*.**
For the bottom-up snake venomics approach the **(A)** RP-HPLC profile was observed at λ = 214 nm, with peak 0 corresponding to the injection peak, and venom fractions further analyzed by **(B)** Coomassie-stained SDS-PAGE under reducing conditions. PAGE line nomenclature is based on RP-HPLC fractions from (A). Labelled bands were cut, subjected to tryptic digestion, and analyzed with LC-MS. For top-down and intact mass profiling the venom profile was observed via the **(C)** total ion current chromatogram by ESI under non-reducing conditions.
